# A Methylation-Phosphorylation Switch Controls EZH2 Stability and Hematopoiesis

**DOI:** 10.1101/2023.02.02.526767

**Authors:** Pengfei Guo, Keshari Rajawasam, Tiffany Trinh, Hong Sun, Hui Zhang

**Author notes:** Co-corresponding Authors: Hong Sun and Hui Zhang, (Hong Sun) (Hui Zhang).

## Abstract

The Polycomb Repressive Complex 2 (PRC2) methylates H3K27 to regulate development and cell fate by transcriptional silencing. Alteration of PRC2 is associated with various cancers. Here, we show that mouse *Lsd1/Kdm1a* deletion causes dramatic reduction of PRC2 proteins, whereas mouse null mutation of *L3mbtl3* or *Dcaf5* results in PRC2 accumulation and increased H3K27 trimethylation. The catalytic subunit of PRC2, EZH2, is mono-methylated at lysine 20 (K20), promoting EZH2 proteolysis by L3MBTL3 and the CLR4^DCAF5^ ubiquitin ligase. LSD1 demethylates the methylated K20 to stabilize EZH2. K20 methylation is inhibited by AKT-mediated phosphorylation of serine 21 in EZH2. Mouse K20R mutants develop hepatosplenomegaly associated with high GFI1B expression, and K20R mutant bone marrows expand hematopoietic stem cells and downstream hematopoietic populations. Our studies reveal that EZH2 is regulated by methylation dependent proteolysis, which is negatively controlled by AKT-mediated S21 phosphorylation to establish a methylation-phosphorylation switch to control the PRC2 activity and hematopoiesis.

## INTRODUCTION

The Polycomb Repressive Complex 2 (PRC2) epigenetically regulates embryonic development and cell fate determination (Chou et al., 2011; Sparmann & van Lohuizen, 2006). The core components of PRC2 include EZH2 (enhancer of Zeste homolog 2), SUZ12 (suppress of Zeste 12), EED (embryonic ectoderm development), and histone binding proteins RbAp48/46. EZH2 acts as the catalytic subunit of PRC2 to trimethylate histone H3 at lysine 27 (H3K27me3) to promote epigenetic gene silencing during development. Mouse *Ezh2*, *Suz12*, or *Eed null* mutant embryo*s* either cease to develop after implantation or initiate but fail to complete gastrulation (Faust et al., 1995; O’Carroll et al., 2001; Pasini et al., 2007). PRC2 represses developmental regulators in mouse embryonic stem cells and deletion of *Ezh2* compromises the self-renewal and differentiation of human embryonic stem cells by de-repressing developmental regulators (Collinson et al., 2016; Shan et al., 2017; Sparmann & van Lohuizen, 2006). PRC2 is required for various other developmental functions including B-lymphoid development, myogenic differentiation, and imprinted X-chromosome inactivation, and loss of PRC2 also exhausts bone marrow hematopoietic stem cells (Chou et al., 2011; Sparmann & van Lohuizen, 2006). The pathological role of EZH2 is highlighted by over-expression or gain-of-function mutations in various cancers (Kim & Roberts, 2016). EZH2 is considered as an important marker for the aggressive stages of prostate and breast malignancies due to its high expression levels in these cancers (Kleer et al., 2003; Varambally et al., 2002). EZH2 also serves as a critical therapeutic target of human malignancies including hematopoietic cancers (Bodor et al., 2013; Chang & Hung, 2012; Kim & Roberts, 2016). Many studies indicate that the post-translational modifications of EZH2 play an important role in cancer development (Li et al., 2020). One of the critical modifications is the phosphorylation of serine 21 (S21) in EZH2 by AKT, activated by the PI3K signaling cascade (Cha et al., 2005). The phosphorylated S21 was reported to inhibit EZH2 methyltransferase activity on H3K27 (Cha et al., 2005). Although AKT-mediated S21 phosphorylation on EZH2 is reported to facilitate tumorigenesis (Kim et al., 2013), the physiological role and regulation of the S21-phosphorylated EZH2 remain largely unclear.

Protein lysine methylation has been extensively investigated in histones to establish the critical roles of mono-, di-, and tri-methylated lysine residues of histones in modulating chromatin structure and gene expression (Greer & Shi, 2012; Zhang et al., 2012). For example, the methylations of Lys 4 (H3K4), Lys 36 (H3K36), Lys 48 (H3K48), and Lys 79 (H3K79) in histone H3 are typically associated with transcriptional gene activation, but the methylations of Lys 9 (H3K9) and Lys 27 (H3K27) on histone H3, or Lys 20 (H4K20) on histone H4 are usually connected to transcriptional silencing (Greer & Shi, 2012). Emerging evidence indicates that many non-histone proteins, such as p53, DNA (cytosine-5)-methyltransferase 1 (DNMT1), NFκB/RelA, ERα, GLI3, SOX2, LIN28A, HIF1α, and E2F1, are mono-methylated on specific lysine residues by SET7 (SET7/9, SET9, SETD7 or KMT7)(Fu et al., 2016; S. K. Kim et al., 2014; Lee et al., 2017; Leng, Yu, et al., 2018; Zhang et al., 2012), originally isolated as a histone methyltransferase that mono-methylates H3K4 (Nishioka et al., 2002; Wang et al., 2001). Our recent studies revealed that specific lysine-methylation on a group of non-histone proteins such as DNMT1, SOX2, SMARCC1, SMARCC2, and E2F1 by SET7 causes the lysine methylation dependent proteolysis of these non-histone proteins (DiPatri et al., 2015; Guo et al., 2022; Zhang et al., 2018; Zhang et al., 2019). We and other also found that the methyl groups on these methylated non-histone proteins are removed by LSD1 (also called KDM1a), initially identified as a histone demethylase that specifically removes methyl groups from the mono- and di-methylated H3K4, but not the trimethylated H3K4, to repress transcription (Guo et al., 2022; Leng, Yu, et al., 2018; Shi et al., 2004; Zhang et al., 2018; Zhang et al., 2012).

In this study, we analyzed the phenotypes of nestin-Cre mediated deletion of mouse *Lsd1* gene. The mouse null mutation of *Lsd1* causes early embryonic lethality (Wang et al., 2009; Wang et al., 2007). Loss of mouse *Lsd1* also profoundly impairs the self-renewal and differentiation of various stem/progenitor cells such as embryonic stem cells and hematopoietic stem cells (Adamo et al., 2011; Saleque et al., 2007; Zhang et al., 2018; Zhang et al., 2019). However, the molecular targets of LSD1 deficiency that cause these pathological defects remain largely unclear. We found that loss of LSD1 affects the levels of PRC2 and our further analyses reveal that the protein stability of EZH2 is regulated by lysine methylation dependent proteolysis.

## RESULTS

### Loss of mouse LSD1 in the mouse brain diminishes the levels of PRC2 proteins

We have bred the floxed *Lsd1* conditional deletion mice (Saleque et al., 2007) with the nestin-Cre transgenic mice (Tronche et al., 1999) to specifically delete *Lsd1* in the central and peripheral nervous system, including neuronal and glial cell precursors. Homozygous loss of *Lsd1* conditional alleles by the nestin-Cre caused the animal lethality immediately after birth on day one (Figure 1A). To determine whether loss of *Lsd1* affects PRC2, we examined the levels of PRC2 proteins in the *Lsd1* mouse mutants. Immunostaining of brain sections from the wildtype and *Lsd1* null mutant animals revealed that the protein level of EZH2 is prominently reduced in the *Lsd1* mutant mice (Figure 1A). Further characterization repeatedly showed that the protein levels of key components of PRC2, including EZH2, SUZ12, and EED, are markedly reduced in the brain extracts of *Lsd1* null mutants, as compared with that of wildtype littermates (Figure 1B). The *Lsd1* deletion induced reduction of these PRC2 protein levels occurred post-transcriptionally, since the PRC2 mRNA levels are comparable between the *Lsd1* null mutants and the wildtype littermates in the brain tissues (Figure 1C). To further examine if the reduction of PRC2 proteins is caused by animal lethality, we established mouse embryonic fibroblasts (MEFs) from the homozygous floxed *Lsd1* conditional (fl/fl) deletion mouse embryos with the actin-Cre-ER (CAGGCre-ER/*Lsd1*fl/fl) by breeding the *Lsd1* conditional deletion mouse strain (Saleque et al., 2007) with a transgenic mouse strain expressing a tamoxifen inducible Cre-ER recombinase under the actin promoter control (CAGGCre-ER)(Hayashi & McMahon, 2002). While the wildtype MEFs normally express substantial levels of PRC2 proteins, induced deletion of *Lsd1* in the CAGGCre-ER/*Lsd1*fl/fl MEFs by addition of 4-hydroxytamoxifen (4-OH-Tam) led to the rapid disappearance of these PRC2 proteins and reduction of H3K27me3 (Figure 1D). Since loss of a single core component of PRC2, such as EZH2 or SUZ12, typically leads to the disassembly of PRC2 and proteolysis of other PRC2 subunits (Collinson et al., 2016; Pasini et al., 2004), we further determined if the protein stability of PRC2 proteins such as EZH2 is dependent on *Lsd1*. We treated the CAGGCre-ER/*Lsd1*fl/fl MEFs with a LSD1 inhibitor, CBB3001, that we previously developed (Guo et al., 2022; Hoang et al., 2018) and found that the protein level of EZH2 is significantly decreased by CBB3001 (Figure 1E). We also used siRNA-mediated silencing of LSD1 in cultured human breast carcinoma T47D cells and cervical carcinoma HeLa cells. We found that LSD1 silencing caused the downregulation of EZH2 protein and the LSD1 silencing-induced EZH2 reduction is reversed by the treatment of 26S proteasome inhibitor, MG132, added in the last 6 hours of the experiment (Figure 1F and 1G)(Guo et al., 2022). Our studies indicate that LSD1 is required to maintain the protein stability of EZH2 to prevent the disassembly of PRC2.

**Figure 1.**
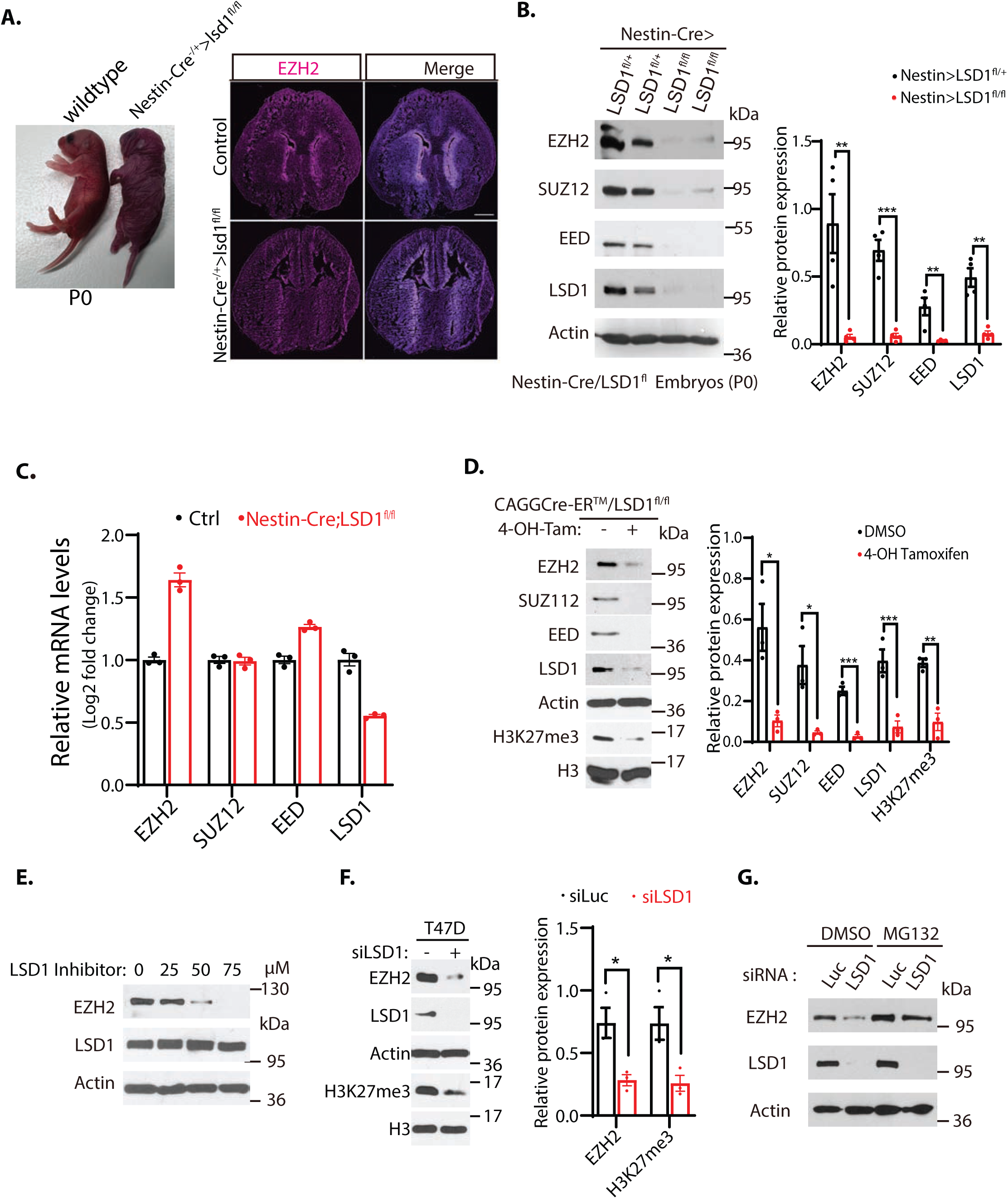
Downregulation of EZH2 in LSD1 null mice. (**A**) Left: Nestin-Cre directed conditional inactivation of mouse *Lsd1* gene causes immediate postnatal death after birth (P0). Right: Immunofluorescence staining of EZH2 from the brain sections of the wildtype and nestin-Cre;LSD1^fl/fl^ mice on P0 day. Scale bars, 500 μm. (**B**) PRC2 protein levels in the brain extracts of P0 Nestin-Cre;LSD1^fl/+^ heterozygous control or nestin-Cre;LSD1^fl/fl^ homozygous *Lsd1* conditional deletion mice were analyzed by Western blotting. (**C**) Reverse-transcriptional quantitative PCR (RT-qPCR) analysis of the mRNA levels of EZH2, SUZ12, EED and LSD1 in the brain of the LSD1^flox/flox^/nestin-Cre mice. The mRNA levels were measured in triplicated by RT-qPCR. Quantifications are represented by bar graph with mean and standard deviation (S.D.) for error bars from three replicate samples and normalized to the control wildtype LSD1^flox/flox^ mice. (**D**) Excision of *Lsd1* by 4-OH-TAM in the LSD^fl/fl^-actin-Cre^ER^ MEFs reduces PRC2 proteins. Embryonic fibroblasts from CAGGCre-ER^TM^/LSD1^fl/fl^ mouse embryos (E13.5) were treated with 4-hydroxytamoxifen (20 μg/ml) for 12 hours to delete *Lsd1* by inducible actin-Cre-ER. (**E**) Wildtype MEFs were treated with various concentrations of LSD1 inhibitor CBB3001 (50 μM) for 20 hours and EZH2 and LSD1 protein levels were analyzed by blotting with indicated antibodies. (**F)** T47D cells were transfected with 50 nM luciferase control or LSD1 siRNAs for 48 hours and the levels of indicated proteins were analyzed. (**G**) H1299 cells were transfected with 50 nM luciferase (Luc) control or LSD1 siRNAs for 48 hours and added 5 μg/ml MG132 for the last 6 hours before lysing the cells for blotting. For (**B**), (**D**) and **(F)**, Significance was indicated as two-tailed, unpaired, *t* test. Values are expressed as the mean ± SEM. \**p*<0.05. \*\**p*<0.01. \*\*\**p*<0.001. Protein molecular weight markers are in kilodalton (kDa).

### Deletion of mouse *L3mbtl3* gene causes the accumulation of EZH2 protein

Since LSD1 serves as a demethylase for the mono- and di-methylated histone H3K4 and several mono-methylated non-histone proteins including DNMT1, E2F1, SOX2, SMARCC1 and SMACC2 (Guo et al., 2022; Leng, Yu, et al., 2018; Zhang et al., 2018; Zhang et al., 2019), we wondered whether PRC2 proteins are regulated by the lysine methylation-dependent proteolysis pathway through L3MBTL3, a methyl lysine reader that binds to mono-methylated DNMT1, SOX2, SMARCC1. Our initial study is to examine and characterize EZH2 and SUZ12 proteins in *L3mbtl3* null mouse embryos (Guo et al., 2022). We repeatedly found that the levels of EZH2 and SUZ12 proteins are significantly increased in the mouse *L3mbtl3* null embryos, which died at E17.5-19.5, as compared to that of the wildtype littermates (Figure 2A). In mouse embryonic fibroblasts isolated from the wildtype or *L3mbtl3* homozygous deletion mutant embryos, the loss of *L3mbtl3* caused the accumulation of EZH2 protein and the increased level of trimethylated H3K27 (Figure 2B). Since the homozygous loss of mouse *L3mbtl3* impairs the maturation of the mouse hematopoietic system to cause anemia and embryonic lethality around E17.5-E19.5 (Arai & Miyazaki, 2005), we employed mouse *L3mbtl3^tm1a(EUCOMM)Hmgu^* embryonic stem cells to generate the conditional floxed *L3mbtl3^fl/fl^* mice by removing the neo-LacZ elements with the Flp recombinase to establish the loxP sites that flank the exon 5 of the *L3mbtl3* allele in the mice (Figure 2-figure supplement 1). To induce the homozygous *L3mbtl3^fl/fl^* conditional deletion in the central nervous system, the *L3mbtl3^fl/fl^* mice were bred with nestin-Cre transgenic mice. We found that homozygous loss of *L3mbtl3* in the brain of the *L3mbtl3^flox/flox^*/nestin-Cre mice survived, but the brain extracts accumulated EZH2 and SUZ12 proteins, as compared with that of *L3mbtl3^flox/+^*/nestin-Cre mice (Figure 2C). Our immunostaining of the brain sections from the wildtype and *L3mbtl3^flox/flox^*/nestin-Cre null mutant animals also revealed that the protein levels of EZH2 and trimethylated H3K27 accumulate in the *L3mbtl3* conditional null mutant brains, as compared to that of the control wildtype littermates (Figure 2D). To further confirm that EZH2 is regulated by LSD1 and L3MBTL3 *in vivo*, the *L3mbtl3* conditional deletion mouse strain were bred with the *Lsd1* conditional deletion strain and the transgenic mouse expressing a tamoxifen inducible Cre-ER recombinase under the actin promoter control (CAGGCre-ER) to generate the conditional knock-out mice of either LSD1 or L3MBTL3 alone, or the double conditional knock-out mice of both LSD1 and L3MBTL3. The mouse embryonic fibroblasts (MEFs) were established from the homozygous CAGGCre-ER/*Lsd1*^fl/fl^ and the double CAGGCre-ER/*Lsd1*^fl/fl^/*L3mbtl3*^fl/fl^ deletion embryos. While these MEFs normally express substantial levels of EZH2 and H3K27me3, induced deletion of *Lsd1* in the CAGGCre-ER/*Lsd1*^fl/fl^ MEFs by addition of 4-hydroxytamoxifen (4-OH-Tam) led to the disappearance of EZH2 protein and the corresponding reduction of H3K27me3 (Figure 2E). However, the induced co-deletion of *Lsd1* and *L3mbtl3* in the CAGGCre-ER/*Lsd1*^fl/fl^/*L3mbtl3*^fl/fl^ MEFs leads to the restoration of the EZH2 and H3K27me3 protein levels in the MEFs. We also used siRNA-mediated silencing of L3MBTL3 in human lung carcinoma H1299 cells with two representative siRNAs (Guo et al., 2022; Leng, Yu, et al., 2018). Our study revealed that while LSD1 silencing reduced EZH2 protein levels, co-silencing of L3MBTL3 and LSD1 stabilized EZH2 protein levels in LSD1 deficient cells (Figure 2F and 2G). Collectively, these studies indicate that loss of LSD1 induced the proteolytic degradation of EZH2 through the L3MBTL3-dependent proteolysis during embryonic development and in cultured cells.

**Figure 2.**
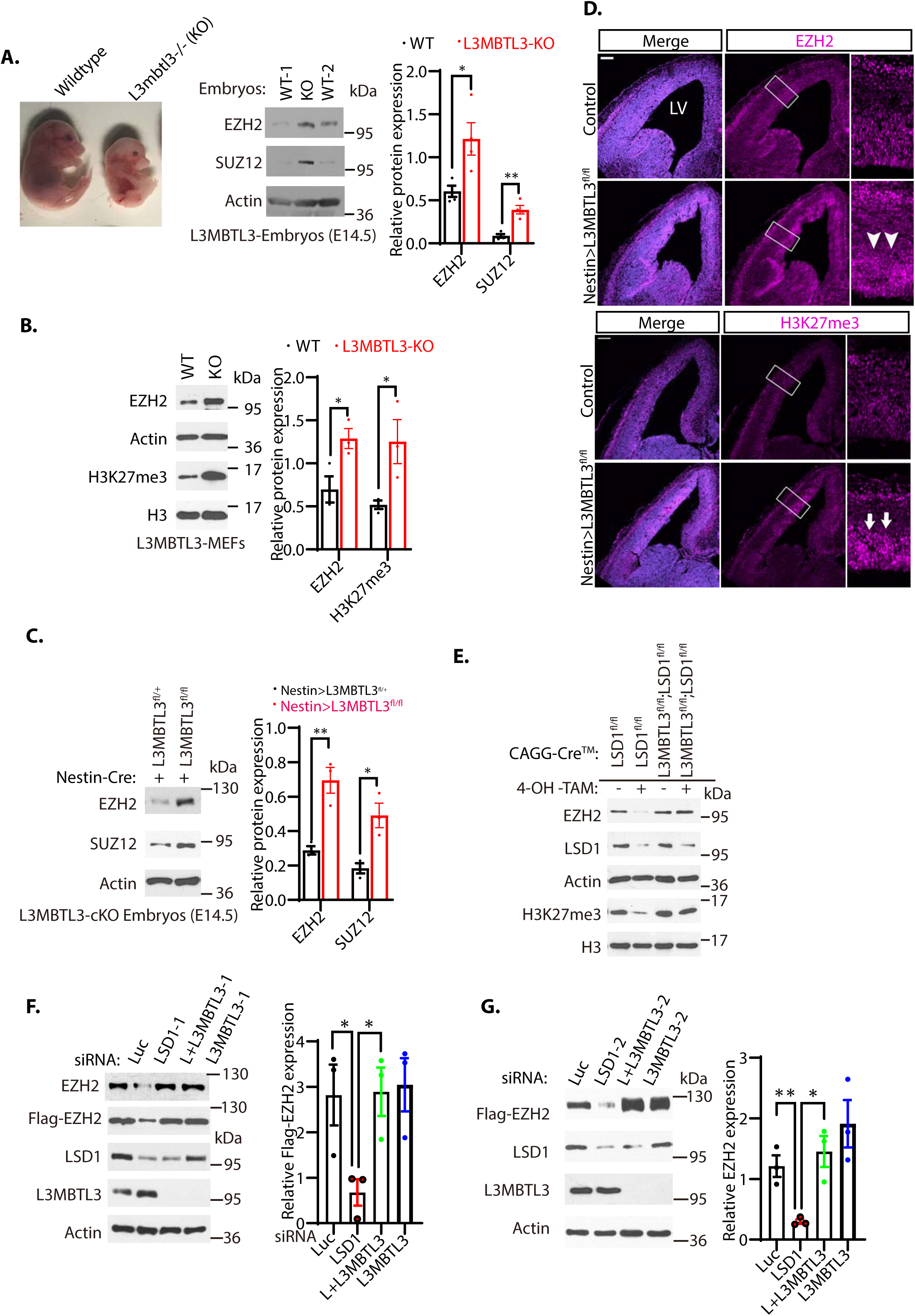
Loss of L3MBTL3 stabilizes EZH2 protein. (**A**) Left: The mouse *L3MBTL3* wildtype (+/+) and *L3MBTL3* null (-/-, KO) mutant embryos on embryonic day 17.5 (E17.5) after breeding. Right: Total lysates from the heads of mouse *L3mbt3 (+/+)* wildtype and *L3mbt3* homozygous deletion (-/-, KO) mutant embryos (equal total proteins) were analyzed by Western blotting with antibodies for the indicated proteins. (**B**) Mouse embryonic fibroblasts from the wildtype and *L3mbtl3* deletion mutant embryos (E13.5) were examined for EZH2 and H3K27me3 proteins by Western blotting. (**C**) Western blot analysis of EZH2 and SUZ12 proteins in the head extracts of nestin-Cre;L3MBTL3^fl/+^ control and nestin-Cre;L3MBTL3^fl/fl^ conditional deletion (cKO) embryos (E15.5) using the indicative antibodies. (**D**) Coronal sections of the developing mouse brain at E15.5 were stained with anti-EZH2 and H3K27me3 antibodies in wildtype control and nestin-Cre;L3MBTL3^fl/fl^ conditional deletion mice. Scale bars, 100 μm. Arrows and arrowheads indicate the regions of EZH2 and H3K27me3 expression, respectively. LV: lateral ventricle. (**E**) Deletion of both LSD1 and L3MBTL3 by 4-OH-TAM restores the protein levels of EZH2 and H3K27me3. MEFs from the nestin-Cre;LSD1^fl/fl^ and CAGGCre-ER^TM^;LSD1^fl/fl^;L3MBTL3^fl/fl^ mouse embryos (E13.5) were treated with 4-hydroxytamoxifen (4-OH-TAM, 20 μg/ml) for 12 hours to delete LSD1 and L3MBTL3. (**F and G**) Silencing of L3MBTL3 re-stabilizes the protein levels of EZH2 in LSD1 deficient cells. The Flag-EZH2 under the retroviral LTR promoter control were ectopically and stably expressed in H1299 cells and the cells were transfected with 50 nM siRNAs of luciferase, LSD1, and two L3MBTL3 siRNAs. The indicated proteins were analyzed by Western blotting. For (**A**-**C**), (**F**), and (**G**), band intensities were quantified and normalized to that of the luciferase or actin control. Significance was indicated as two-tailed, unpaired, *t* test. Values are expressed as the mean ± SEM. \**p*<0.05. \*\**p*<0.01. \*\*\**p*<0.001.

### Deletion of mouse *Dcaf5* leads to the accumulation of EZH2 and H3K27me3

Our previous studies have shown that L3MBTL3 recruits CRL4^DCAF5^ ubiquitin E3 ligase complex to target substrates, such as DNMT1, SOX2, and SMARCC1, for ubiquitin dependent proteolysis (Guo et al., 2022; Leng, Yu, et al., 2018; Zhang et al., 2019). To determine whether DCAF5, a substrate specific subunit of the CRL4 ubiquitin E3 ligase complex (Leng, Yu, et al., 2018), is involved in regulating the levels of EZH2 and H3K27me3, we employed the CRISPR-Cas9 gene editing system to delete the exon 4 of the mouse *Dcaf5* allele to establish the *Dcaf5* deletion mutant mouse line with a stop codon to the remaining downstream read frame of the *Dcaf5* allele (Figure 3A and 3B)(S. Kim et al., 2014; Kleinstiver et al., 2016; Slaymaker et al., 2016). We found that homozygous mutation of the *Dcaf5* alleles also caused significant elevation of EZH2 and H3K27me3 protein levels (Figure 3C). Immunostaining of the embryonic brain sections revealed that loss of *Dcaf5* leads to increased levels of EZH2 and H3K27me3 staining, as compared with that of control wildtype littermates (Figure 3D). Consistent with these animal studies, siRNA-mediated co-silencing of DCAF5 and LSD1 in H1299 cells with two representative siRNAs stabilized EZH2 protein in LSD1 deficient cells (Figure 3E and 3F). These results indicate that the CRL4^DCAF5^ ubiquitin ligase complex is involved in the proteolytic degradation of EZH2 to regulate the H3K27me3 levels during mouse development and in cultured cancer cells.

**Figure 3.**
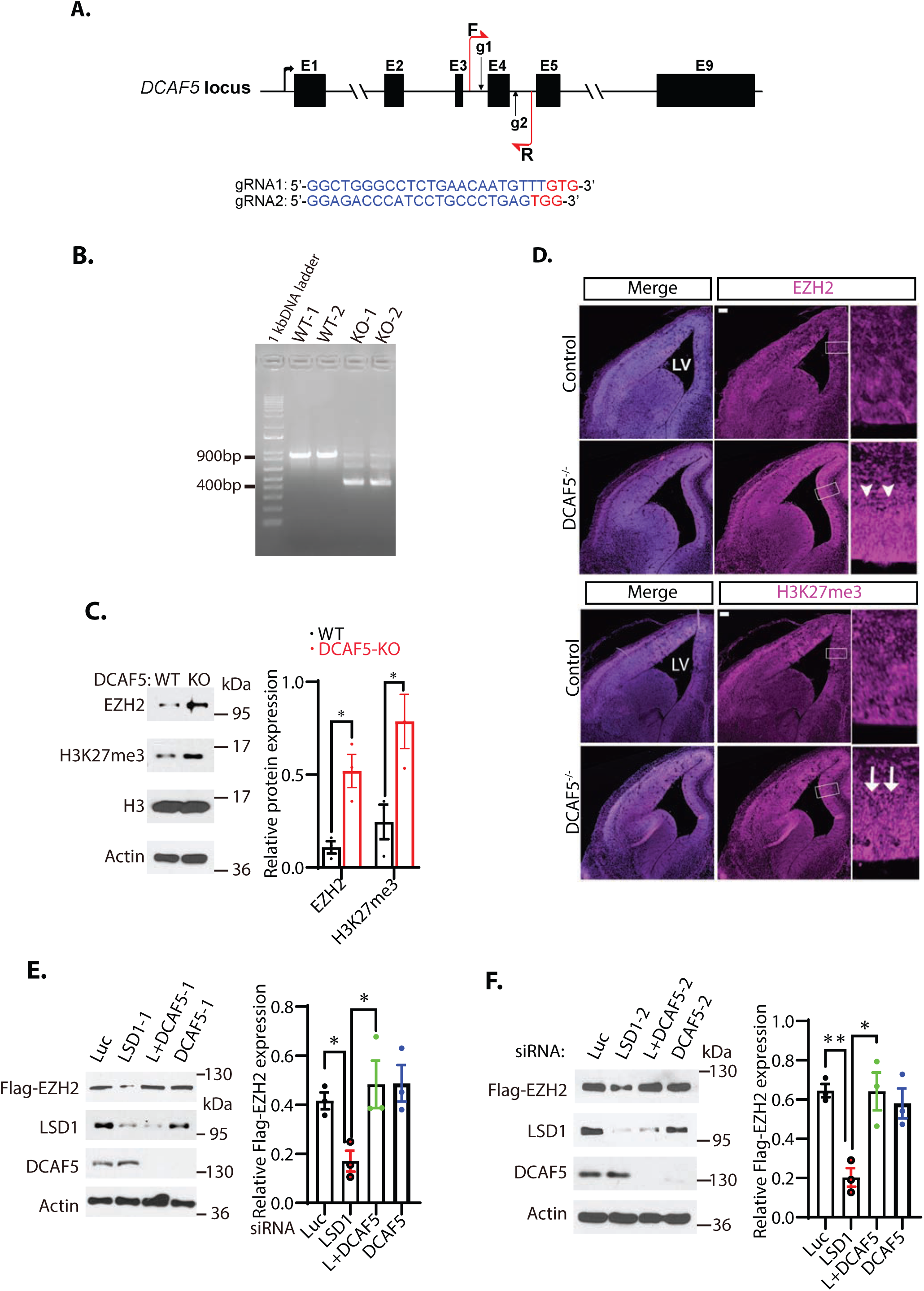
Loss of DCAF5 stabilizes EZH2 protein. **A.** The strategy to delete the exon 4 of mouse *Dcaf5* gene by CRISPR-Cas9 gene edition with two guide RNAs (gRNAs). (**B**) Genome typing of the wildtype (WT) and *Dcaf5* knock-out (KO) mice by PCR. (**C**) Western blot analysis of EZH2 and H3K27me3 proteins in the brains of the wildtype control and homozygous *Dcaf5* deletion mice using the indicative antibodies. (**D**) Accumulation of EZH2 and H3K27me3 proteins in mouse *Dcaf5* deleted embryonic brains. Immunostainings of anti-EZH2 and H3K27me3 in the coronal sections of the mouse embryonic brains of the wildtype control and *Dcaf5* at E15.5. Scale bars, 100 μm. Arrows and arrowheads indicate the expression regions of EZH2 and H3K27me3, respectively. Boxed regions are enlarged on the right panels. LV: lateral ventricle. **(E)** and **(F**) Silencing of DCAF5 re-stabilizes the protein levels of EZH2 in LSD1 deficient cells. H1299 cells expressing stably expressed Flag-EZH2 were transfected with 50 nM siRNAs of luciferase, LSD1, and two DCAF5 siRNAs. The indicated proteins were analyzed by Western blotting. Band intensities in (**C**), (**E**), and (**F**) were quantified and normalized to that of the histone H3 or luciferase control. Significance was indicated as two-tailed, unpaired, *t* test. Values are expressed as the mean ± SEM. \**p*<0.05. \*\**p*<0.01.

### The protein stability of EZH2 is regulated by lysine methylation

Our mouse genetic evidence indicates that the protein stability of EZH2 is regulated by LSD1, L3MBTL3, and DCAF5, which are involved in regulating the proteolytic degradation of lysine methylated protein substrates such as DNMT1, SOX2, SMARCC1 and SMARCC2 (Leng, Saxena, et al., 2018; Leng, Yu, et al., 2018; Zhang et al., 2019). To further investigate EZH2 regulation, we examined and found that EZH2 contains a conserved lysine residue, Lys20 (K20), in the putative SET7 methylation consensus motif (R/K-S/T/V-K) that is very similar to that of the Lys42 (K42) methylation degron motif in SOX2 (Figure 4A)(Zhang et al., 2018; Zhang et al., 2019). To test whether the K20 residue in EZH2 serves as a putative substrate for LSD1, we synthesized the monomethylated K20 peptide and its unmethylated cognate peptide derived from EZH2 (Figure 4-figure supplement 1). We mixed and incubated the monomethylated K20 peptide with purified GST-LSD1 and found that the recombinant GST-LSD1 protein can effectively remove the methyl group of the monomethylated K20 peptide (Figure 4C), indicating that the monomethylated K20 in EZH2 serves as a direct substrate of LSD1.

**Figure 4.**
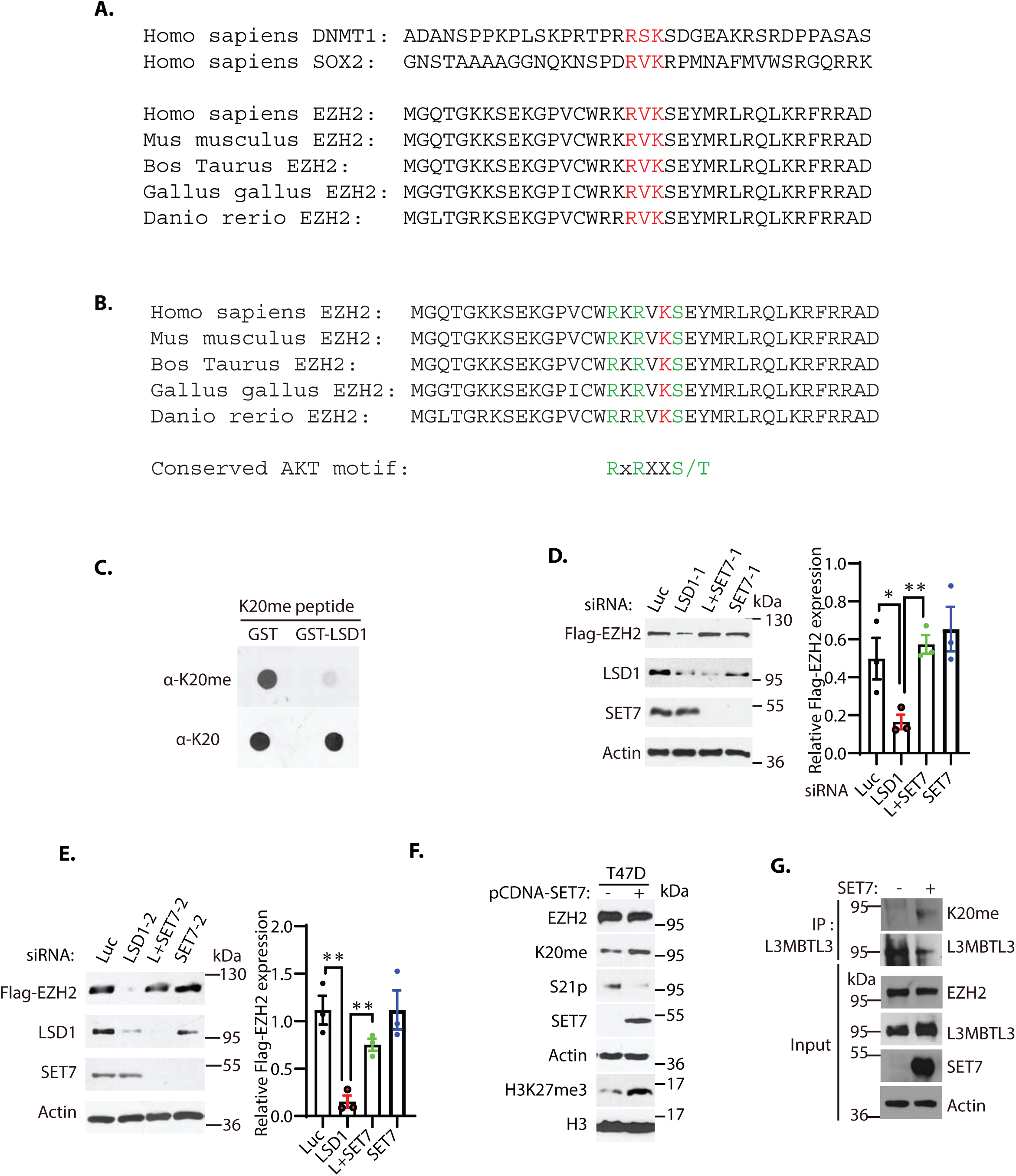
The K20 residue in EZH2 is methylated by SET7. **A.** EZH2 contains a conserved lysine residue, Lys 20 (K20), within the consensus lysine residues (K*) of the H3K4-like R/K-S/T/V-K* methylation motifs methylated by SET7. (**B**) K20 (red) is next to serine 21 (S21) that is phosphorylated by AKT in in of EZH2. The critical amino acid residues (RXRXXS) in the AKT phosphorylation consensus motif are labeled green. **(C)** LSD1 demethylates the methylated K20 peptide. Purified 1 μg GST control or GST-LSD1 proteins were incubated with 100 ng of mono-methylated K20 peptides for 4 hours at room temperature and the resulting peptides and input methylated peptides were spotted onto nitrocellulose membrane and blotted with affinity purified anti-methylated K20 and anti-K20 antibodies as indicated. (**D** and **E**) Silencing of SET7 re-stabilizes the protein levels of EZH2 in LSD1 deficient cells. H1299 cells expressing stably expressed Flag-EZH2 were transfected with 50 nM siRNAs of luciferase, LSD1, and two independent SET7 siRNAs. The indicated proteins were analyzed by Western blotting. The protein bands were quantified and normalized to that of the luciferase control. Significance was indicated as two-tailed, unpaired, *t* test. Values are expressed as the mean ± SEM. \**p*<0.05. \*\**p*<0.01. (**F**) T47D cells were transfected with control vector (pcDNA3) or SET7 expression construct for 48 hours and protein extracts were prepared. Proteins were detected by Western blotting with indicative antibodies. **(G)** The K20-methylated EZH2 preferentially binds to L3MBTL3. The 293 cells were transfected with control vector (pcDNA3) or SET7 expression construct for 48 hours, and interactions between L3MBTL3 and EZH2-K20me were analyzed by co-immunoprecipitation and Western blotting analyses.

We next assessed whether SET7 methyltransferase is involved in the regulation of EZH2 proteolysis. We found that while silencing of LSD1 reduced the protein level of ectopically and stably expressed Flag-tagged EZH2 in H1299 cells, co-silencing of SET7 with two siRNAs and LSD1 effectively re-stabilized Flag-EZH2 protein in LSD1 deficient cells (Figure 4D and 4E). To effectively detect the methylated K20 in EZH2, we generated and affinity purified a specific anti-mono-methylated K20 (K20me) peptide antibody for EZH2, which only recognized the mono-methylated K20me peptide (Figure 4-figure supplement 1), but not the unmethylated cognate K20 peptide. Using this affinity purified K20me antibody, we examined K20 methylation of EZH2 in T47D cells (Figure 4F). We found that ectopic expression of SET7 led to substantially increased K20 methylation of EZH2 and H3K27me3 proteins (Figure 4F) and the K20-methylated EZH2 fraction is enriched in the L3MBTL3 co-immunoprecipitation with EZH2 (Figure 4G). These studies indicate that SET7 is capable of methylating K20 in EZH2, and that the methylated K20 binds to L3MBTL3. However, SET7 expression also led to a substantial reduction of the phosphorylated serine 21 (S21) in EZH2, a site previously reported to be phosphorylated by the active AKT activity (Figure 4F)(Cha et al., 2005). We noticed that the K20 residue is immediately next to serine 21 (S21) in EZH2 (Figure 4A, 4B, and Figure 4-figure supplement 1). Previous studies have shown that S21 of EZH2 is phosphorylated by the activated AKT to suppress the activity of EZH2 to trimethylated H3K27, but the S21 phosphorylation does not alter EZH2 protein stability (Cha et al., 2005). Our results indicate that SET7 catalyzes the mono-methylation of K20 and also cause the decrease of the phosphorylation of S21 in EZH2, suggesting that the methylation of K20 may negatively regulate the phosphorylation of S21 of EZH2. This notion was also supported by our finding of the inverse correlation between the levels of K20 methylation and S21 phosphorylation in EZH2 during mouse embryonic development (Figure 5A and 5B). Our studies revealed that the methylated K20 levels of EZH2 gradually increased during mouse embryogenesis from E14.5, E18.5, to the first postnatal day after birth (P0), whereas this K20me increase was concurrently accompanied by the gradual reduction of the S21-phosphorylated form of EZH2, phosphorylated AKT, and the total EZH2 protein levels during the indicated developmental stages, even though the total AKT protein levels remain high throughout the process (Figure 5A and 5B).

**Figure 5.**
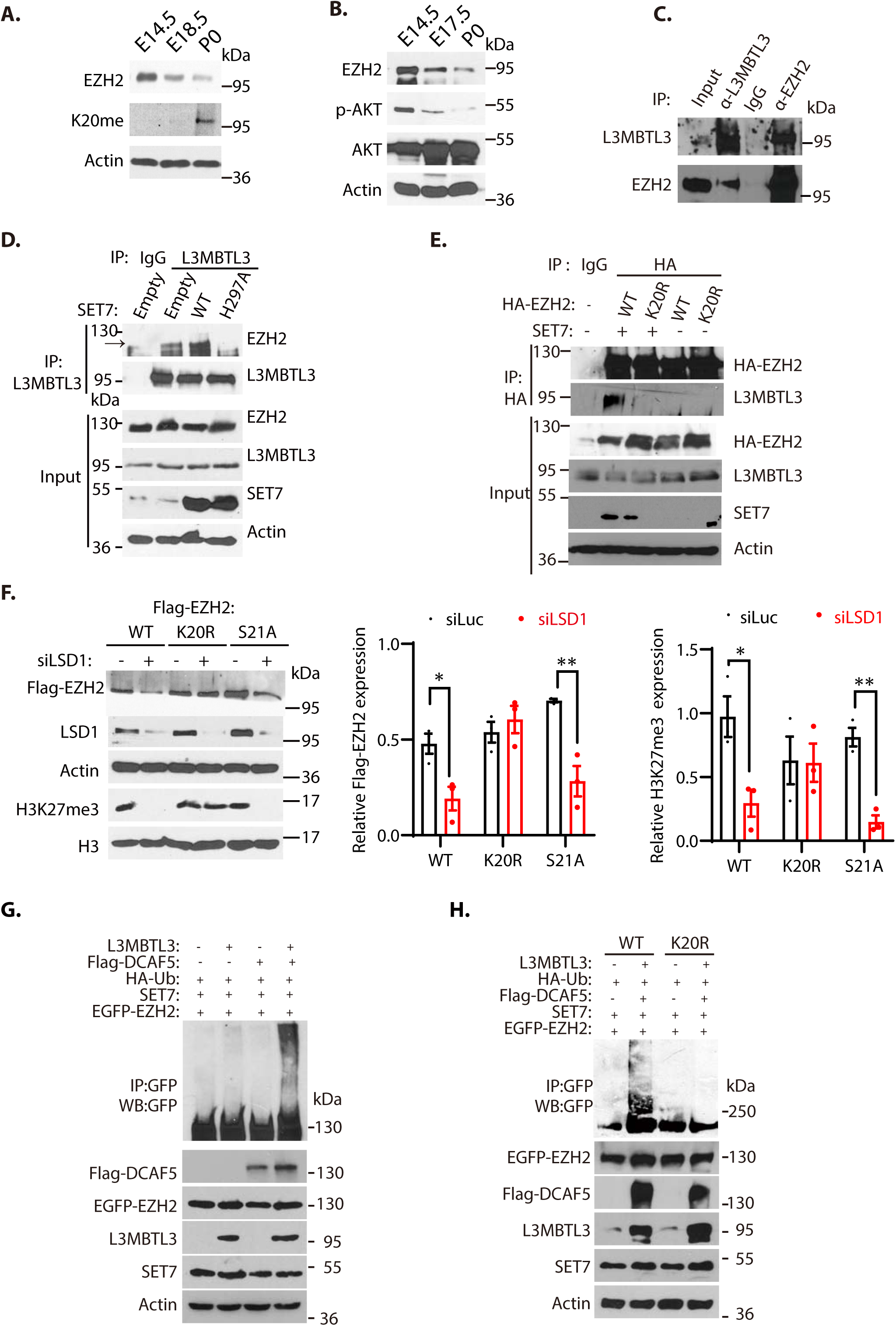
L3MBTL3 and DCAF5 target methylated K20 to promote EZH2 proteolysis. (**A** and **B**) Mouse embryo lysates at the indicated embryonic days were prepared and the indicated proteins were examined with respective antibodies. **(C)** Endogenous L3MBTL3 interacts with EZH2. Lysates were extracted from mouse embryos (E14.5) and the interaction between L3MBTL3 and EZH2 were analyzed by co-immunoprecipitation and blotted with respective antibodies. Input: 1/10 of the lysates for Western blotting. (**D**) SET7 stimulates the interaction between L3MBTL3 and EZH2. 293 cells were transfected with expression vector of SET7 wildtype, SET7 H297A mutant, or the empty vector for 48 hours. Cell lysates were immunoprecipitated by anti-L3MBTL3 antibodies and blotted with antibodies against EZH2 and L3MBTL3. (**E**) The K20R mutant does not interact with L3MBTL3. HA-tagged EZH2 or the HA-K20R mutant expressing constructs were co-transfected with SET7 or empty vector into 293 cells. Cell lysates were immunoprecipitated with the anti-HA antibody and the blots were immunoblotted with anti-L3MBTL3 and anti-HA antibodies. (**F**) The Flag-tagged EZH2 wildtype, K20R, or S21A mutant were stably expressed in G401 cells. The cells were then transfected with 50 nM siRNAs of luciferase or LSD1 for 48 hours. The protein levels of Flag-EZH2, H3K27me3, and indicated other proteins were analyzed by immunoblotting. The protein bands were quantified and normalized to that of the luciferase control. Significance was indicated as two-tailed, unpaired, *t* test. Values are expressed as the mean ± SEM. \**p*<0.05. \*\**p*<0.01. (**G** and **H**) The EGFP-tagged wildtype EZH2 or K20R mutant were co-transfected into 293 cells together with vectors expressing HA-tagged ubiquitin (HA-Ub) and SET7 in the presence or absence of L3MBTL3 and DCAF5 expressing constructs as indicated. Proteins were immunoprecipitated with anti-GFP antibodies and Western blotted with anti-GFP and other antibodies against indicated proteins.

In mouse embryonic extracts, our co-immunoprecipitation analysis revealed that L3MBTL3 and EZH2 interact (Figure 5C); and the interaction between L3MBTL3 and EZH2 was promoted in the presence of an active SET7 methyltransferase, but not an inactive SET7 mutant in which the critical histidine 297 is converted to alanine (H297A, Figure 5D). In addition, the SET7-promoted binding of EZH2 to L3MBTL3 is dependent on the presence of K20 in EZH2, as the conversion of K20 to arginine (K20R) in EZH2 abolished the EZH2-L3MBTL3 interaction (Figure 5E). We also found that while the wildtype EZH2 is reduced by LSD1 silencing, the K20R mutant of EZH2 is resistant to the loss of LSD1 (Figure 5F), although the S21A mutation of EZH2 that converts S21 to alanine still remains to be sensitive to LSD1 silencing, indicating that K20 is still methylated (Figure 5F). Furthermore, co-expression of SET7, L3MBTL3, and DCAF5 are sufficient to promote EZH2 polyubiquitination (Figure 5G), but the K20R mutant of EZH2 under the same conditions failed to be polyubiquitinated (Figure 5H). These studies collectively indicate that K20 of EZH2 is methylated by SET7, and that L3MBTL3 and CRL4^DCAF5^ recognize and target the K20-methylated EZH2 protein for ubiquitination dependent proteolysis.

### The methylation of K20 and phosphorylation of S21 are mutually exclusive in EZH2

Our studies indicate that the levels of K20-methylation and S21-phosphorylation of EZH2 have an inverse relationship during mouse embryonic development (Figure 5A and 5B). To further examine whether loss of LSD1 affects the protein stability of wildtype EZH2, K20R, and S21A mutant EZH2, human rhabdoid tumor G401 cells stably expressing the HA-tagged wildtype, K20R-, and S21A-EZH2 were transfected with LSD1 siRNA and then treated with protein synthesis inhibitor, cycloheximide, to block translational initiation to measure protein decay rates (Guo et al., 2022). We found that the half-lives of EZH2 proteins were reduced after LSD1 silencing in cells expressing the wildtype and S21A mutant EZH2, but the EZH2R mutant is quite resistant to LSD1 silencing (Figure 6-figure supplement 1). However, the initial S21 phosphorylation studies did not reveal that the block S21-phosphorylation by AKT inhibition resulted in the reduction of EZH2 protein stability in T47D cells (Cha et al., 2005). Indeed, we found that treatment of T47D cells with MK2206, an AKT inhibitor, inhibited AKT-mediated S21 phosphorylation of EZH2 and increased levels of H3K27me3 due to the reactivation of EZH2 methyltransferase activity, but the total EZH2 protein levels were not significantly altered (Figure 6A), consistent with the previous report (Cha et al., 2005). However, we found that ectopic expression of SET7 in T47D cells caused the increased K20 methylation in EZH2 and elevated levels of H3K27me3, accompanied with the reduced S21 phosphorylation of EZH2 (Figure 4F), suggesting that the methylation of K20 by SET7 inhibits the phosphorylation of S21 in EZH2 to activate the H3K27 methyltransferase activity of EZH2 in T47D cells. To further investigate the regulation of EZH2 protein stability by AKT phosphorylation of S21, we examined the response of primary wildtype MEFs to MK2206 by measuring EZH2 protein levels. We found that MK2206 reduced the levels of EZH2 protein, the phosphorylated AKT, phosphorylated S21 in EZH2, and consequent H3K27me3 levels, concurrent with increased levels of the K20 methylated EZH2 protein (Figure 6B). Notably, we treated H1299 cells that stably express Flag-tagged EZH2 under the retroviral LTR promoter control with various doses of MK2206. We found that increasing concentrations of MK2206 caused the downregulation of both Flag-EZH2 and endogenous EZH2 proteins, associated with the reduced levels of H3K27me3 (Figure 6C). These results are consistent with our observation in MEFs and SET7-expressing T47D cells (Figure 4F, 5A and 5B), and show that reduction of S21 phosphorylation in MEFs and H1299 cells facilitates the methylation of K20 by SET7, leading to EZH2 proteolysis and consequently reducing the trimethylated H3K27 levels. These studies indicate that EZH2 is regulated by a methylation-phosphorylation switch of EZH2 in MEFs and H1299 cells but the K20-methylation dependent degradation of EZH2 is not detectable after the MK2206 treatment in T47D cells. We tried to determine the cause of differential response of T47D cells to AKT inhibition and we found that ectopic and stable expression of L3MBTL3 in T47D cells conferred the EZH2 proteolytic response to MK2206. As shown in Figure 6D, we ectopically expressed the HA-tagged L3MBTL3 in T47D cells and then treated the T47D cells with MK2206. Our analysis revealed that inhibition of AKT leads to the reduced level of EZH2 protein and consequent downregulation of H3K27me3 levels in the T47D cells that ectopically expressing L3MBTL3, similar to that of MEFs and H1299 cells (Figure 6D). We reasoned that the EZH2 protein levels in T47D cells did not significantly respond to AKT inhibition is likely caused by the weak L3MBTL3-dependent proteolysis activity of EZH2 in this cell line and we tried to further investigate this possibility by using the *L3mbtl3* null MEFs that is deficient in targeting the K20-methylation dependent proteolysis of EZH2 (Figure 2B). Our studies revealed that MK2206 treatment caused increased levels of H3K27me3 in the *L3mbtl3* null MEFs due to the removal of S21-phosphorylation mediated inhibition on EZH2 and PRC2, whereas the AKT inhibition did not cause any further changes in EZH2 protein levels because of *L3mbtl3* deletion, a result similar to the response of T47D cells to MK2206 (Figure 6A and 6E). These studies are consistent with our hypothesis that T47D cells may have reduced activities in L3MBTL3 and CRL4^DCAF5^ ubiquitin E3 ligase activities to target the K20-methylated EZH2 for proteolysis. Our results are also consistent with other reports showing that AKT inhibition by MK2206 reduced EZH2 protein stability in several cancer cells (Riquelme et al., 2016), indicating the methylation-phosphorylation switch of EZH2 exists in many cells but T47D is defective in this pathway. Since *L3mbtl3* is mutated in medulloblastoma and is further implicated in other pathological disorders such as multiple sclerosis, insulin resistance, prostate cancer and breast cancer (Andlauer et al., 2016; Bonasio et al., 2010; Kar et al., 2016; Lotta et al., 2017; Northcott et al., 2009), we found that the levels of L3MBTL3 protein are differentially expressed in various cancer cell lines and that T47D cells express relatively low levels of L3MBTL3 protein in the cell lines we analyzed (Figure 6-figure supplement 2). However, it is likely that the lysine methylation dependent proteolysis is affected at various levels including the altered L3MBTL3 protein levels in cancer cells to control EZH2 protein levels. Further investigation is required to clarify the alterations of L3MBTL3 in various cancer cells.

**Figure 6.**
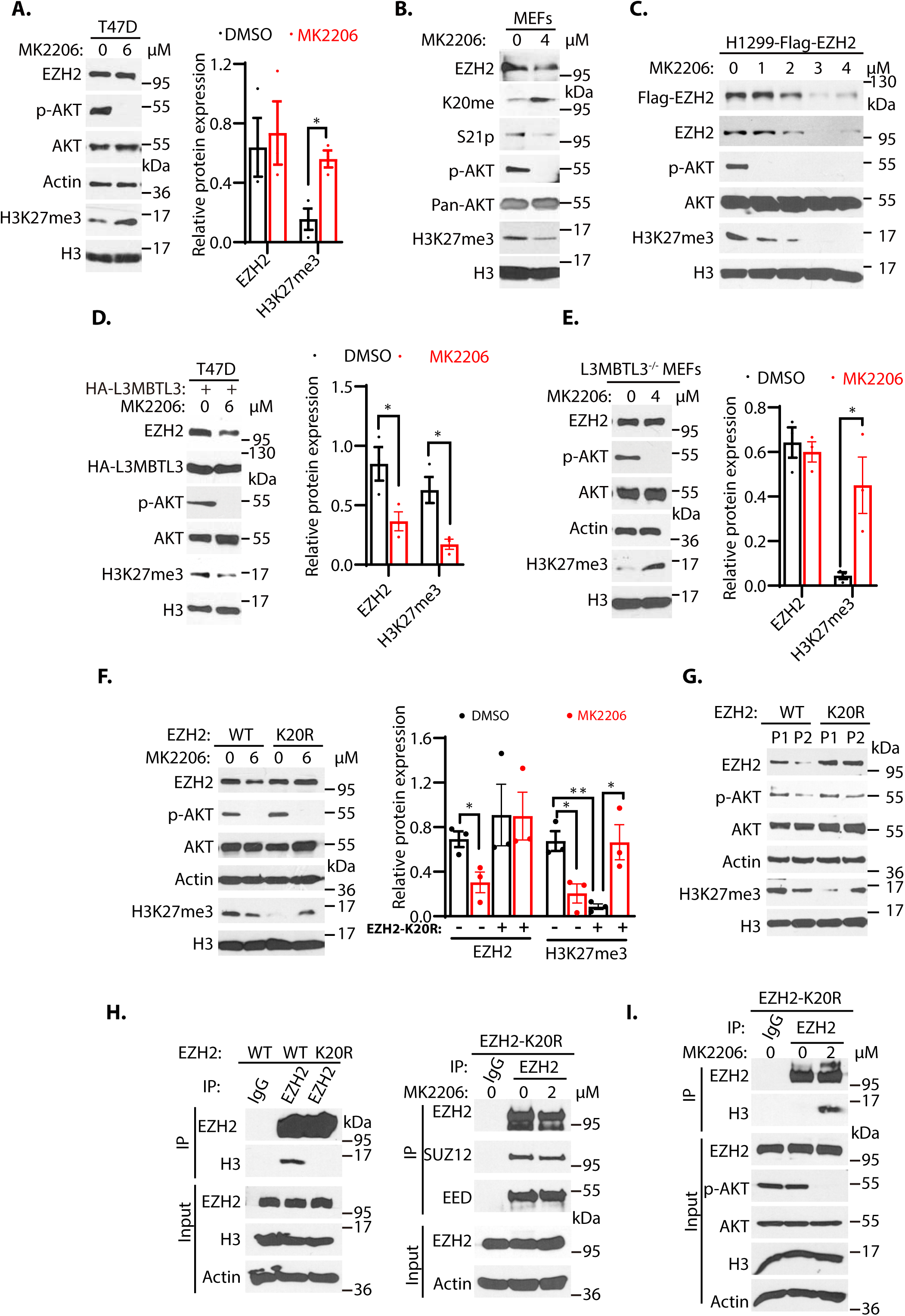
The methylation-phosphorylation switch regulates the stability of EZH2. (**A-C**) T47D cells (**A**), mouse embryonic fibroblasts (MEFs, **B**), or H1299 cells stably expressing Flag-EZH2 (**C**) were treated with AKT inhibitor MK2206 (6 μM) or dimethyl sulfoxide (control, DMSO) as indicated for 4 hours. Protein lysates were blotted with indicated antibodies. (**D**) T47D cells were transfected with the HA-L3MBTL3 expression construct for 48 hours and the cells were then treated with or without 6 μM MK2206 for 4 hours. The indicated proteins were blotted with specific antibodies. (**E**) Primary L3mbtl3 (-/-) MEFs were cultured and the cells were treated with or without 4 μM MK2206 for 4 hours. The indicated proteins were blotted with specific antibodies. (**F**) Primary MEFs were obtained from the EZH2 wildtype and homozygous K20R mutant embryos (E14.5). They were treated with DMSO or MK2206 for 4 hours. Lysates were prepared and proteins were blotted with indicated antibodies. (**G**) Primary MEFs were obtained from the EZH2 wildtype and homozygous K20R mutant mouse embryos (E14.5) and cultured as passage 1 (P1). They were passaged by splitting 1/3 and cultured as passage 2 (P2). Lysates were prepared and proteins were blotted with indicated antibodies. (**H**) Lysates from primary EZH2 wildtype and K20R mutant MEFs were immunoprecipitated with antibodies against EZH2. The proteins were blotted with anti-EZH2 and histone H3 antibodies. (**I**) Primary EZH2 K20R mutant MEFs were treated with or without MK2206 (2 μM) for 4 hours. The lysates were immunoprecipitated by anti-EZH2 antibodies. Proteins were blotted with anti-SUZ12 and anti-EED antibodies. For (**A**), (**D**), (**E**), and (**F**), protein band intensities were quantified and normalized to that of histone H3 or actin control. Significance was indicated as two-tailed, unpaired, *t* test. Values are expressed as the mean ± SEM. \**p*<0.05. \*\**p*<0.01.

### Establishment and characterization of K20R mutant mice

To systematically determine the effect of the K20 methylation on EZH2 and its relationship to S21 phosphorylation, we employed the CRISPR-Cas9 gene editing technique to change the nucleotides AA for codon 20 of the mouse *Ezh2* gene to GG, resulted in converting K20 to arginine (K20R) in the mouse (S. Kim et al., 2014; Kleinstiver et al., 2016; Slaymaker et al., 2016). The targeted K20R knock-in allele (K20R^ki^ or K20R) was confirmed by direct DNA sequencing of the heterozygous and homozygous K20R^ki^ mouse strains (Figure 6-figure supplement 3). The primary homozygous K20R MEFs and wildtype control MEFs were isolated and cultured from the homozygous K20R and wildtype mouse embryos. These MEFs were treated with MK2206 and the wildtype and K20R of EZH2 proteins were examined. Our results revealed that while the EZH2 and H3K27me3 proteins in wildtype MEFs were reduced by MK2206, the K20R mutant of EZH2 was resistant to AKT inhibition (Figure 6F). However, the control treated primary K20R MEFs showed lower H3K27me3 levels than that of the wildtype control MEFs, likely because the S21 of the K20R protein can still be phosphorylated by the AKT activity to inhibit PRC2 catalytic activity for the H3K27me3 levels. Notably, MK2206 treatment induced the increased levels of H3K27me3 in the K20R MEFs (Figure 6F), similar to the response of T47D cells and the *L3mbtl3* null MEFs to MK2206 (Figure 6A and 6E). Previous studies reported that the protein levels of EZH2 decline during the *in vitro* passages of primary MEFs (Bracken et al., 2007). We tried to determine whether the EZH2 protein level decline is dependent on K20 methylation by culturing both K20R and the corresponding wildtype MEFs. Our studies indicate that while the *in vitro* cell passage from passage 1 to passage 2 induces the down-regulation of wildtype EZH2 protein and H3K27me3 levels in the primary MEFs, the K20R mutant protein is resistant to the passage-dependent decline (Fig. 6G). In addition, the H3K27me3 level increased in the passage 2 of the K20R mutant MEFs from the lower levels in the passage 1 of the K20R mutant MEFs, likely caused by the downregulation of phosphorylated and active AKT in the passage 2 of the K20R mutant MEFs (Figure 6G). We tried to examine the changes of S21 phosphorylation during the cell passages but the S21 phosphorylation signal is too low to be detected by our anti-S21 phosphorylation antibodies. Nevertheless, our results indicate that the K20R mutant stabilizes EZH2 protein to increase H3K27 trimethylation by PRC2, and that loss of AKT phosphorylation on S21 of EZH2 promotes K20 methylation and EZH2 proteolysis, whereas loss of K20 methylation conversely facilitates S21 phosphorylation and stabilizes EZH2 protein.

Since S21 phosphorylation was reported to reduce EZH2 interaction with histone H3 (Cha et al., 2005), we analyzed the interaction between EZH2 and histone H3 by immuno-co-precipitation. Our results showed that the K20R mutant of EZH2 has less association with histone H3 than the wildtype EZH2 protein in MEFs, although their interaction with SUZ12 or EED did not change (Figure 6H). It is likely that the K20R mutation in EZH2 may facilitate the phosphorylation of S21 by AKT to block the interaction between EZH2 and histone H3. Indeed, we found that AKT inhibition enhanced the interaction between the K20R mutant and histone H3 (Fig. 6I). It has been shown that the Notch1 intracellular domain (NICD) increases the cytoplasmic EZH2 levels during early megakaryopoiesis due to AKT-dependent phosphorylation of EZH2 (Roy et al., 2012). We examined the distribution of the K20R mutant in MEFs. Immunofluorescent staining revealed that while the wildtype EZH2 protein was nuclear with little cytoplasmic presence, a small fraction of the K20R protein was found to localize to the cytoplasm (Figure 6-figure supplement 4A), although the majority of EZH2 still remains in the nucleus. We also performed cell fractionation studies and our biochemical analysis indicate that a fraction of the K20R mutant protein is localized in the cytoplasmic fraction, but the majority of EZH2R mutant protein is still in the nucleus (Figure 6-figure supplement 4B). Thus, our results indicate that the K20R mutation abolishes the methylation of K20 in EZH2 and consequently promotes S21 phosphorylation by AKT to prevent EZH2 to interact with histone H3 and that the K20R mutation promotes the presence of a small fraction of EZH2 in the cytoplasm, likely due to the increased S21-phosphorylation in the K20R mutant of EZH2.

### The K20R mutation causes the expansion of bone marrow hematopoietic populations

To determine the *in vivo* functional deficit of the K20R mutation of EZH2, we bred the heterozygous K20R^ki/wt^ mice to generate the homozygous K20R knock-in mutant (K20R^ki/ki^ or K20R) mice. The homozygous K20R progeny mice are viable with normal Mendelian genetic ratios. We found that the livers and spleens of the 8-month-old homozygous K20R mutant mice are enlarged (hepatosplenomegaly), as compared with that of the wildtype littermates (Figure 7A and 7B). In addition, the total cell numbers of the bone marrow (BM) in 5-month-old mice were significantly increased in the K20R mice than in that of wildtype animals (Figure 7C). We next investigated the potential role of the K20R mutation of EZH2 in regulating the primitive hematopoietic subpopulations in the bone marrow. The homozygous K20R mutant mice contained a significantly higher absolute number of the Lin^-^Sca1^+^c-Kit^+^ (LSK) cells, which represent multipotent stem cells that are CD48^+^, CD71^+^, and enriched for CD150^+^ capable of regenerating bone marrow from irradiated mice, and LK cells, which are c-Kit^low^ and enriched for Sca-1^+^ progenitor cells, as compared with that of the wildtype control mice (Figure 7D-F)(Adolfsson et al., 2001; Beguelin et al., 2013). Analysis of the LK cells of the K20R mice revealed an obvious expansion in the numbers of granulocyte-monocyte progenitor (GMP), common myeloid progenitor (CMP), and myeloid erythroid progenitor (MEP) populations (Figure 7G). Moreover, our flow cytometric analysis revealed that the bone marrow of the K20R mutant mice showed an increased proportion of mature myeloid cells (Gr1^+^/Mac1^+^ and Ly6G) (Figure 7H and 7I), whereas the absolute numbers of B and T lymphocytes remained similar to that of the wildtype animals (Figure 7-figure supplement 1A-C). Our quantitative reverse transcription-polymerase chain reaction (RT-PCR) analysis further showed that the homozygous K20R mutant bone marrow cells displayed significantly increased RNA expression levels of transcriptional regulators for the hematopoietic stem cell (HSC) activities, specification, and self-renewal, including Runx1, GATA2, Etv6, cFOS, IKAROS, and GFI1 (Figure 7J and Table S1)(Deneault et al., 2009; Hock et al., 2004; North et al., 1999; Pereira et al., 2013; Ross et al., 2012; Tsai et al., 1994). These results indicate that the K20R mutant mice promote the expansion of both hematopoietic stem cells and the myeloid lineage in the bone marrow.

**Figure 7.**
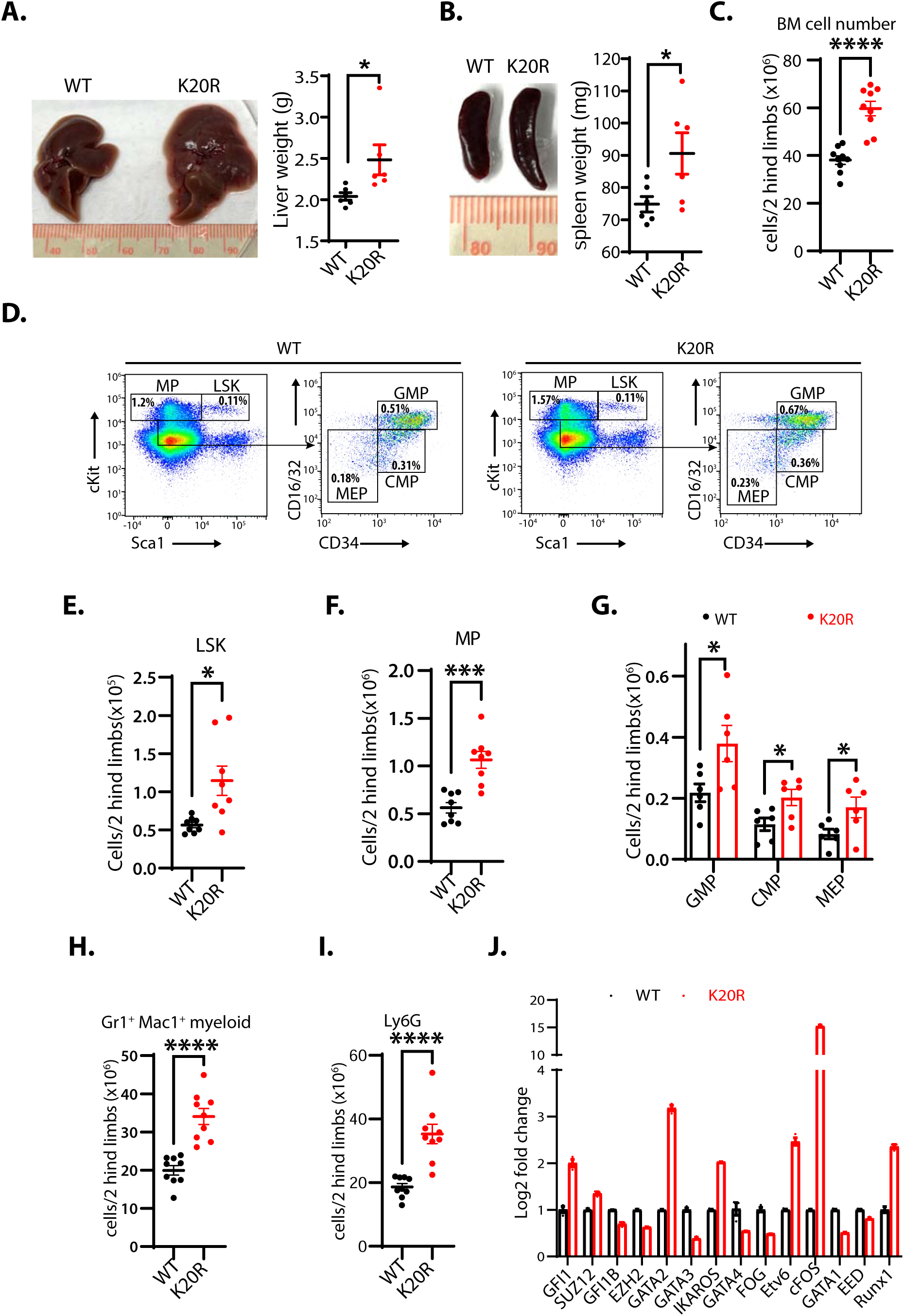
K20R mutation causes hepatosplenomegaly and expansion of hematopoietic populations. (**A** and **B**) The enlarged liver or spleen from the homozygous K20R mutant mice, as compared to that of the EZH2 wildtype mice (8 months old). The weights of livers or spleens from the K20R and wildtype EZH2 mice (7∼10-month-old) were measured and plotted. Values are means ± SEM (N=6, *p<0.05). (**C**) Number of total cells in bone marrows harvested from 5 months old K20R and wildtype EZH2 mice. Values are means ± SEM (N=9, ****p<0.0001). (**D-F**) Representative flow cytometric (FACS) profiles of bone marrow cells from the K20R and EZH2 wildtype mice. Flow cytometry plots were gated on the Lin^-^cKit^+^ (myeloid progenitors) subpopulation that is subclassified into CMP, GMP, and MEP based on Lin^-^cKit^+^ CD16/32 and CD34 expression. The number of immature cells [Lin^-^cKit^+^Scal1^+^(**E**), Lin^-^cKit^+^ (**F**), CMP, GMP, and MEP in (**G**)], and differentiated cells [Mac1^+^Gr1^+^ myeloid (**H**) and Ly6.6G myeloid (**I**) in bone marrow samples harvested from the EZH2 wildtype and K20R mutant mice. Values are means ± SEM (n=6∼9). Significance was indicated as a two-tailed, unpaired, *t-test. *p* <0.05.\*\*\**p*<0.001. ****p<0.0001. (**J**) The quantitative RT PCR (qRT-PCR) analysis shows mRNA expression levels of indicated genes in bone marrow samples harvested from the EZH2 wildtype and K20R mutant mice.

### Phenotypic characterization of the K20R of EZH2 mutant mice

Many studies have shown that EZH2 overexpression is correlated with human hematological malignancies and EZH2 has been established as a critical target for hematological cancers (Abd Al Kader et al., 2013; van Galen et al., 2007; Yan et al., 2013). Since the K20R mutant mice exhibit enlarged spleen and liver (Figure 7A and 7B), our hematoxylin and eosin (HE) staining of the splenic sections from 8-month-old K20R mutant mice revealed the prominent hemosiderin-laden macrophages in the red pulp and mild to moderate reactive hyperplasia in the follicles of the white pulp (Figure 8A). The K20R mutation also caused mild to moderate diffuse cytoplasmic vacuolation in the mouse liver sections (Figure 8A). Furthermore, increased protein levels of EZH2 and H3K27me3 were observed in the spleen of adult K20R mutant mice (Figure 8C), associated with the increased Ki67 immunostaining of splenic sections, indicating increased numbers of proliferating cells in K20R mutant animals (Figure 8B). These findings suggest that the single amino acid substitution of lysine-to-arginine substitution, K20R, in EZH2 modulates stabilized EZH2 protein and induces mild to moderate reactive hyperplasia in animals. To test whether the phenotypes of the K20R mutation are associated with altered transcriptional factors in the spleen, we performed quantitative RT-PCR analysis of the spleens from the wildtype and K20R mutant mice (Figure 8D). Different from the expanded bone marrow cells in the K20R mutant, the quantitative RT-PCR analysis of the K20R spleen cells showed significantly increased expression of GFI1B, an essential proto-oncogenic transcriptional regulator necessary for development and differentiation of erythroid and megakaryocytic lineages (Pereira et al., 2013; Ross et al., 2012). We also assessed and compared the protein levels of GFI1B in the K20R and the wildtype mouse spleens. Increased GFI1B protein levels were detected in the K20R mutant spleens by immunohistological staining with anti-GFI1B antibodies (Figure 8B). The increased GFI1B protein in the K20R mutant spleens were also confirmed by Western blotting analyses of 3- or 7-month-old K20R mutant mice (Figure 8E). Our studies are consistent with a model by which K20 of EZH2 protein is methylated by SET7 to recruit L3MBTL3 and the CRL4^DCAF5^ ubiquitin ligase complex to target the EZH2 protein for ubiquitin dependent degradation, and that LSD1 serves as a demethylase to remove the methyl group from the methylated EZH2 to prevent EZH2 degradation to preserve the integrity of the PRC2 complex (Figure 8F). Importantly, the K20 methylation dependent proteolysis of EZH2 is negatively regulated by the AKT-mediated phosphorylation on S21, which acts to inhibit the H3K27 methyltransferase activity of PRC2 (Figure 8F).

**Figure 8.**
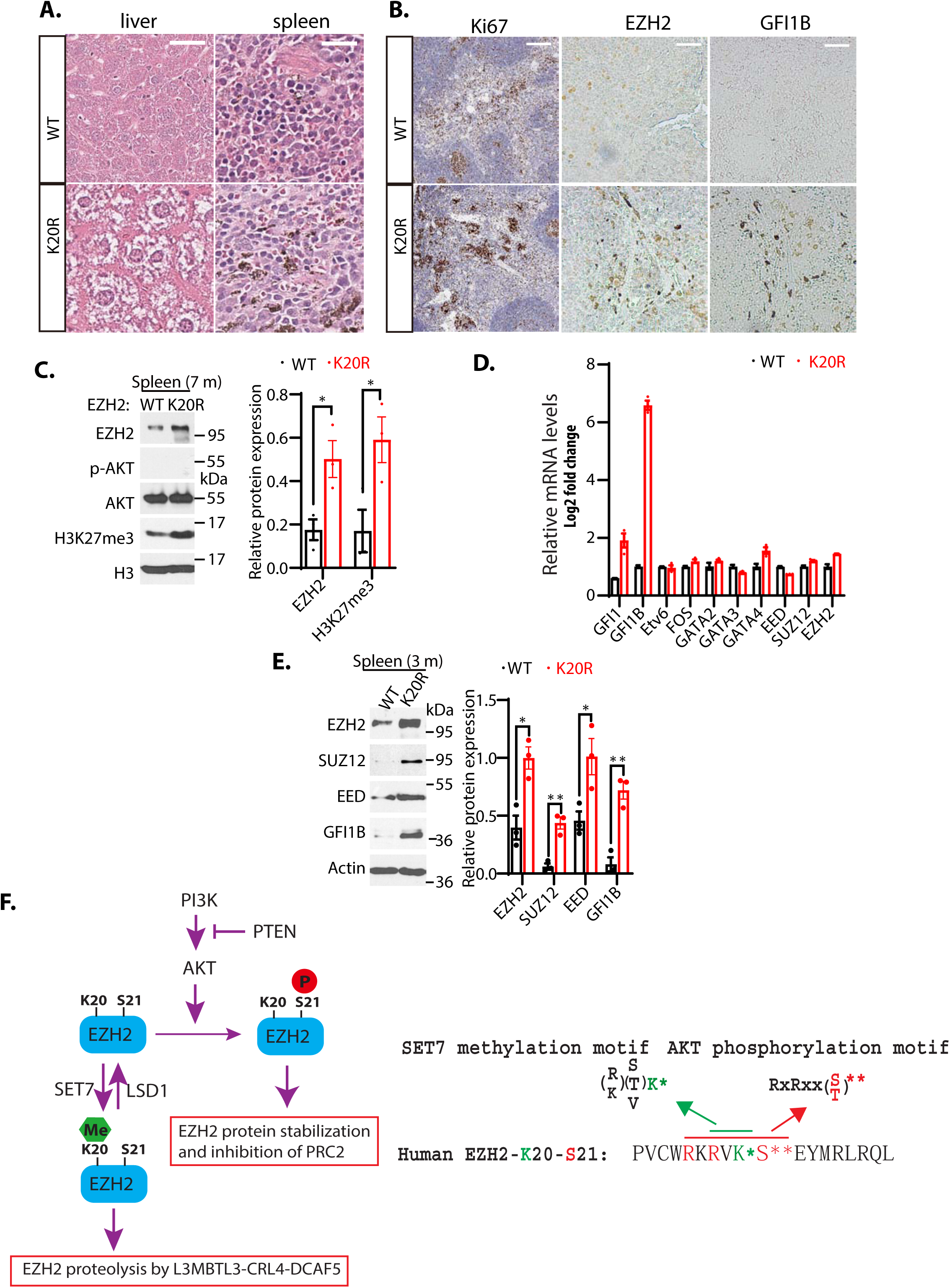
K20R mutation induces hyperplasia in the mouse spleen. (**A**) Hematoxylin and eosin (H&E) histological staining of liver and spleen sections from the K20R and EZH2 wildtype mice (8 months old). Scale bar: 200 μm. **(B)** The anti-Ki67 and anti-GFI1B immunostaining of spleen sections from the K20R and EZH2 wildtype mice (8 months old). Scale bar: 50 mm. (**C**) Left: protein lysates were extracted from the spleen of the K20R and EZH2 wildtype mice and blotted with indicated antibodies. Band intensities were quantified and normalized to that of histone H3 control. Significance was indicated as two-tailed, unpaired, *t* test. Values are expressed as the mean ± SEM. \**p*<0.05. (**D**) RNA levels of indicated hematopoietic regulatory genes were measured by quantitative RT-PCR (RT-qPCR) from the spleens of the EZH2 wildtype and K20R mice (4 months old). The mRNA levels were measured in triplicated by RT-qPCR. (**E**) Protein lysates were extracted from the spleens of the EZH2 wildtype and K20R mice and detected with anti-EZH2, SUZ12, EED, and GFI1B antibodies, using the actin antibody as a control. The protein band intensity values are means ± SEM (N=3). Significance is indicated as a two-tailed, unpaired, *t-test. *p* <0.05.\*\**p*<0.01. (**F**) Model: Left panel: The lysine residue 20 (K20) of EZH2 is methylated by SET7 methyltransferase and the level of methylated K20 is reversibly removed by LSD1 demethylase. L3MBTL3 preferentially binds to the methylated K20 in EZH2 to recruit the CRL4^DCAF5^ ubiquitin E3 ligase complex to target the methylated EZH2 for ubiquitin-dependent proteolysis. Right panel: The K20 methylation is negatively regulated by the phosphorylation of serine 21 (S21) by the PI3K activated AKT. Conversely, the S21 phosphorylation is mutually exclusive to the methylation of K20, resulted in the methylation-phosphorylation switch to control the activity and proteolysis of EZH2 for H3K27 trimethylation.

## Discussion

The alterations of EZH2 and PRC2 are frequently associated with a wide variety of human cancers including hematopoietic malignancies and EZH2 inhibitor, tazemetostat, has been approved for the treatment of follicular lymphoma (Duan et al., 2020). However, how the levels of EZH2 and PRC2 are regulated remain largely unclear. We have previously shown that EED in PRC2 interacts with CRL4 but the significance of this interaction remains unclear (Higa et al., 2006). In this report, we found that the protein stability of EZH2 is dynamically regulated by a novel lysine methylation dependent proteolysis involving the activities of SET7, LSD1, L3MBTL3, and the CRL4^DCAF5^ ubiquitin ligase complex. Our studies revealed that K20 of EZH2 is monomethylated by SET7 methyltransferase and the methylated K20 serves as a substrate of LSD1 demethylase. The methylated K20 is recognized by specific methyl lysine reader L3MBTL3 to promote EZH2 for ubiquitin dependent proteolysis by the CRL4^DCAF5^ ubiquitin E3 ligase complex, resulting the disassembly of the PRC2 complex and reduction of H3K27 trimethylation in animals, MEFs, and cancer cells. Interestingly, the methylation of K20 is prevented by the AKT-dependent phosphorylation of S21 in EZH2. As the active PRC2 complex contains the unphosphorylated EZH2 at S21, which is methylated at K20 to be targeted for proteolysis, further investigation is required to determine how the methylation/phosphorylation switch occurs during normal development and how this regulation is altered in various human diseases including cancers.

## Materials and Methods

### Cells

Human lung carcinoma H1299, cervical carcinoma HeLa, embryonic kidney 293, colon cancer HCT116, rhabdoid tumor G401, breast cancer T47D, teratocarcinoma PA-1, squamous cell lung carcinoma H520, and mouse teratoma F9 cells were purchased from the American Type Culture Collection (ATCC) and cultured in RPMI-1640, McCoy’s 5a, Eagle’s Minimum Essential Medium, or DMEM medium with 10% FBS and 1% antibiotics as described (Guo et al., 2022; Leng, Yu, et al., 2018; Zhang et al., 2013). Mouse embryonic fibroblasts (MEFs) were generated from wildtype and *L3mbtl3* deletion mutant mouse embryos, or the CAGGCre-ER^TM^/LSD1^fl/fl^ and CAGGCre-ER^TM^/LSD1^fl/f^/L3MBTL3^fl/fl^, wildtype and EZH2-K20R knock-in mouse embryos (E12.5-E13.5), as described previously (Guo et al., 2022). For stable expression, human EZH2 wildtype and the K20R or S21A mutant of human EZH2 were cloned into the retroviral pMSCV-Puro vector containing 3xFlag-3xHA epitope (Addgene) and the recombinant retroviruses were packaged in 293 cells (Guo et al., 2022; Leng, Yu, et al., 2018). Viral infected H1299 and G401 cells were selected by puromycin resistance as described before (Guo et al., 2022; Leng, Yu, et al., 2018).

### Peptide synthesis and preparation of methylated peptides

The monomethylated K20 (KSEKGPVCWRKRVK(me1)SEYMRLRQLKRFRRAD) and cognate unmethylated peptides of EZH2 were synthesized at ABI Scientific (Guo et al., 2022). The monomethylated K20 peptide was used to raise rabbit polyclonal antibodies after coupling the peptide to keyhole limpet hemocyanin (KLH)(Guo et al., 2022). Affinity purification of methylated peptide antibodies were conducted as described (Guo et al., 2022; Leng, Yu, et al., 2018). The unmethylated and monomethylated K20 peptides were immobilized to Sulfolink-coupled-resins (Thermo Fisher) by covalently cross-linking with the cysteine residues at the end of the peptides to the resin (Leng, Yu, et al., 2018). The anti-monomethylated K20 peptide sera (5 ml each) were diluted in 1:1 in PBS and first passed through the unmethylated K20 peptide columns (1 ml) for three times to deplete anti-K20 peptide antibodies (Guo et al., 2022). The unbound flow-through antibody fractions were then loaded onto the monomethylated K20 peptide column (0.5 ml), washed, and the bound antibody fractions were eluted by 5 ml of 100 mM glycine, pH2.5. The eluted antibodies (0.5 ml/fraction) were immediately neutralized by adding 100 μl of 2M Tris, pH8.5, and tested for specificity towards the monomethylated K20 peptide but not to the unmethylated K420 peptide (Guo et al., 2022; Leng, Yu, et al., 2018; Zhang et al., 2019). Human LSD1 were cloned into pGEXKG vectors and purified by GSH-Sepharose (GE Healthcare). For demethylation reaction, purified 1 μg of control GST control or GST-LSD1 proteins were incubated with 100 ng of the unmethylated or monomethylated K20 peptides for 4 hours at room temperature, and the resulting peptides were blotted onto nitrocellulose membrane (Guo et al., 2022; Leng, Yu, et al., 2018). The demethylated peptides were detected by immuno-blotting with affinity purified anti-monomethylated K20 antibodies.

### Antibodies and immunological analysis

Anti-LSD1 (A300-215A), L3MBTL3 (A302-852), SUZ12 (A302-407A), and SET7 (A301-747A) antibodies were purchased from Fortis Life Sciences. Anti-EED (ab236292) was purchased from Abcam. Phospho-EZH2 (Ser21) antibody (AF3822) was purchased from Affinity Biosciences. Anti-EZH2 (5246), anti-EED (85322), anti-GFI1B (5849), and anti-H3K27me3 (9733) were from Cell Signaling Technology. Actin (Sc-1616) antibody was purchased from Santa Cruz Biotechnologies. Anti-Flag, HA, and GFP antibodies were purchased from Sigma. Rabbit anti-L3MBTL3 and affinity purified anti-DCAF5 antibodies were also produced in the laboratory as previously described (Leng, Yu, et al., 2018). For direct Western blotting, cells were lysed in the 1 X SDS sample buffer (4% SDS, 100 mM Tris, pH6.8, and 20% glycerol), quantified by protein assay dye (Bio-Rad), and equalized by total proteins (Leng, Yu, et al., 2018). For immunoprecipitation (IP), cells were lysed with a NP40-containing lysis buffer (0.5% NP40, 50 mM Tris, pH 7.5, 150 mM NaCl, and protease inhibitor cocktails)(Leng, Yu, et al., 2018). About 500 μg of lysates and 1μg antibody were used for each IP assay. The antigen–antibody complexes were pulled down by 30 μl Protein A-Sepharose (GE Healthcare) and specific proteins were detected by the Western blotting analysis, using secondary goat anti-mouse HRP (Jackson Immuno Research, 115-035-008) and goat–anti-rabbit antibodies (Jackson Immuno Research, 111-035-008), or Protein A HRP (GE Healthcare, NA9120V), all at 1:2500 dilutions (Leng, Yu, et al., 2018; Zhang et al., 2019).

### Transfection and siRNAs

Oligofectamine was used for siRNA silencing in HeLa, H1299, HCT116, G401, or 293 cells, whereas Lipofectamine 2000 was used for transient transfection as described previously (Leng, Yu, et al., 2018; Zhang et al., 2019; Zhang et al., 2013). Typically, 50 nM of each siRNA or their combinations were transfected into target cells for 48 hours and cells were directly lysed in the 1X SDS lysis sample buffer (Leng, Yu, et al., 2018). For verification of the silencing effects of various target proteins, usually 2 or 3 independent siRNAs were designed to examine the knockdown efficiency and the consequences of knockdown on target proteins (Guo et al., 2022; Leng, Yu, et al., 2018; Zhang et al., 2019). The siRNAs for human genes are: LSD1: GGAAGAAGAUAGUGAAAAC; LSD1-2: UGAAAACUCAGGAAGAUU; DCAF5-1: CUGCAGAAACCUCUACAA; DCAF5-2: ATCACCAACTTCTGACATA; L3MBTL3-1: GATGCAGATTCTCCTGATA; L3MBTL3-2: GGTACCAACTGCTCAAGAA; SET7-1: GGGCAGTATAAAGATAACA; SET7-2 SMART pool: CAACUGCAUCUACGAUAU, CCUGGACGAUGACGGAUUA, GGAGUGUGCUGGAUAUAUU, and CAAACUGG-CUACCCUUAUG (Guo et al., 2022; Leng, Yu, et al., 2018; Zhang et al., 2019). All siRNAs were synthesized from Horizon Discovery.

### Animals

The LSD1^fl/+^ conditional mutant (B6.129-Kdm1a tm1.1Sho/J, stock No: 023969), transgenic actin-Cre-ER (CAGGCre-ER^TM^, B6.Cg-Tg(CAG cre/Esr1*)5Amc/J, stock No: 004682), and transgenic nestin-Cre (B6.Cg-Tg(Nes-cre)1Kln/J, stock No: 003771) mouse strains were obtained from Jackson Laboratory (Guo et al., 2022). The *L3mbt3* deletion mutant (*MBT-1*-/+, B6;129-L3mbtl3tm1Tmiy) mouse strain was previously described (Arai & Miyazaki, 2005; Guo et al., 2022; Leng, Yu, et al., 2018). The *Dcaf5* deletion mutant mouse strain was produced with gRNA1: CTAGTTAGGTACAATAGGGC and gRNA2: TATTCCTCTGCGACCACTCA, flanking the exon4 of the mouse *Dcaf5* locus with the altered read-frame in the downstream of protein sequence, in Centre for Phenogenomics (Toronto, Canada)(S. Kim et al., 2014; Kleinstiver et al., 2016; Slaymaker et al., 2016). The null *Dcaf5* mutant mice are alive and initially bred with wildtype mice for more than 10 generations to ensure the knock-out effects. The EZH2 K20R knock-in mice were produced with the gRNA: ACACGCTTCCGCCAACAAAC and the repair template of a single-strand oligonucleotide with the nucleotide changes encoding c.59_60AA>GG required for the K20R change of mouse *Ezh2* at Phenogenomics (S. Kim et al., 2014; Kleinstiver et al., 2016; Slaymaker et al., 2016). The mouse *L3MBTL3^tm1a(EUCOMM)Hmgu^* embryonic stem cells containing the verified conditional LoxP sites flanking the exon 5 of *L3mbtl3* were obtained from European Mouse Mutant Cell Repository (EuMMCR) and the *L3mbtl3tm1a(EUCOMM)Hmgu* mice were produced at University of California, Davis/KOMP Repository. The *L3mbtl3tm1a(EUCOMM)Hmgu* mice were bred with the FLPo-10 mouse strain (B6.Cg-Tg(Pgk1-flpo)10Sykr/J, Stock No: 011065) from Jackson Laboratory to delete the LacZ and Neo cassettes to establish the *L3mbtl3* ^fl/+^ conditional mutant mice (Wu et al., 2009). All the mutant mice were DNA sequenced and verified. All animal experiments including breeding, housing, genotyping, and sample collection were conducted in accordance with the animal protocols approved by the institutional Animal Use and Care Committee (IACUC) and complied with all relevant ethical regulations at University of Nevada, Las Vegas. All procedures were conducted according to the National Institutes of Health (NIH) Guide for Care and Use of Laboratory Animals. The UNLV IACUC is an AAALAC approved facility and meets the NIH Guide for the Care and Use of Animals.

### Animal phenotype analysis

For mouse embryos analyses, usually 3 pairs of the heterozygous (-/+) male and female mice (10-12 weeks old) in three cages, each with 1 male and 1 female, were bred in the late afternoon and the breeding plugs were examined in the female mice in next morning (Guo et al., 2022). The positive plugs were count as embryonic day 1 (E1) and the pregnant female mice between E14-E17.5 were euthanized by the primary method of CO_2_ asphyxiation, followed by cervical dislocation (secondary method), as approved by the institutional IACUC committee (Guo et al., 2022). Usually a single pregnant female mouse produced about 6-8 embryos, which segregated at the Mendelian inheritance ratio, usually with 1-2 *L3MBTL3* (-/-) or *DCAF5* (-/-), 1-2 wildtype and 3-4 heterozygous *L3MBTL3* (-/+) or *DCAF5* (-/+) embryos. The *L3MBTL3* null embryos between E17.5-19.5 usually died and became disintegrated so they were excluded from protein analyses (Arai & Miyazaki, 2005; Guo et al., 2022; Leng, Yu, et al., 2018). For the analysis of PRC2 proteins in LSD1^fl/fl^/nestin-Cre mice. Usually 3-4 pairs of the LSD1^flox/flox^ male and LSD1^flox/+^/nestin-Cre mice female mice (10-12 weeks old) were bred (Guo et al., 2022). The animals were collected immediate after birth to avoid any delay in sample analysis. The brains of the mice were dissected for protein or immunostaining analysis. For immunostaining, embryos or dissected brains were fixed in 4% paraformaldehyde (PFA) at 4°C overnight and embedded in Optimal cutting temperature compound (O.C.T) according to standard procedures (Christopher et al., 2017; Guo et al., 2022). Sections (10 µm thick, coronal) were stained with specific antibodies and counter-stained with 4′,6-diamidino-2-phenylindole (DAPI)(Guo et al., 2022). Images were acquired with the Nikon A1Rsi Confocal LSM. The sample size was chosen on the basis of our experience on *L3MBTL3, DCAF5, EZH2-K20R,* or *Lsd1* mutant mice and on cultured cells in order to detect the EZH2 and H3K27me3 proteins for differences of at least 50% between the wildtype and mutant groups (Guo et al., 2022). For analysis of *EZH2-K20R* mice, the heterozygous *EZH2-K20R* mice were bred to obtain the wildtype, EZH2-K20R heterozygous, and EZH2-K20R homozygous mice, as the K20R homozygous mutants survive. In the experimental analyses for examination of proteins, the investigators were unaware of the genotypes of the experimental embryos. The investigators also randomly analyzed the wildtype, heterozygous and homozygous knockdown embryos. For the analysis of proteins and DNA from embryos, the experimental procedures for embryo isolation were approved by the UNLV Institutional Animal Use and Care Committee (IACUC). The embryos from the euthanized pregnant female mice or dissected brains, spleens, or livers from the conditional knockout mice, washed with PBS, and lysed in the NP40 lysis buffer (Guo et al., 2022). For the nuclear and cytosolic fractions were separated by centrifugation. Genomic DNA was isolated from nuclear pellets by Zymo genomic DNA-tissue prep kit and quantified. Proteins in the cytosolic suppernant of the lysates were quantified by protein assay dye (Bio-Rad), equalized, and boiled for 15 minutes after addition of 1% SDS and 5% beta-mercaptoethanol to the lysates. Proteins were resolved in protein gel and analyzed by Western blotting (Guo et al., 2022).

### Flow cytometry

Flow cytometry analyses were performed using SONY SH800 high-speed multilaser flow cytometer and cell sorter with the FlowJo^TM^ software in the Core Facility of Nevada Institute of Personalized Medicine. Single-cell suspensions were harvested from bone marrow and lysed with the ACK buffer (ThermoFisher). For mature cells, cells were analysed with directly conjugated anti-mouse antibodies (Biolegend): Ly-6G-PE (1A8), Gr-1-PE (RB6-8C5), CD11b/Mac1-APC (M1/70), CD4-PE/Cyanine7 (GK1.5), CD8a-APC (53-6-7), and B220-PE (RA3-6B2)(Akashi et al., 2000; Arai & Miyazaki, 2005; Traver et al., 2001). For immature cells, depletion of lineage cells were labeled with a cocktail consisting of biotinylated antibodies (Biolegend): Gr-1 (RB6-8C5), TER-119, CD3e (145-2C11), and CD11b (M1/70), conjugated to the EasySep^TM^ Mouse Streptavidin RapidSpheres^TM^ (Cat: #19860A, STEMCELL Technologies), and separated on the EasySep^TM^ Magnet (Cat:#18000) according to the accompanying protocol from STEMCELL Technologies (Akashi et al., 2000; Arai & Miyazaki, 2005; Traver et al., 2001). For inmature cells, directly conjugated antibodies used were as follows: streptavidin-FITC (Cat: 405201), c-Kit- PE/Cyanine7 (2B8), Sca-1-PE/Dazzle 594 (D7), CD16/CD32-PE (2.4G2), and CD34 Alexa Fluor 647 (RAM34). Dead cells were stained with Zombie Green^TM^ (Cat: 423111)(Akashi et al., 2000; Arai & Miyazaki, 2005; Traver et al., 2001). All these antibodies were used at 1:150.

### RNA extraction and qRT-PCR analysis

RNA was extracted from the heads of mouse embryos or neonatal mice using Trizol regent (ThermoFisher) according to the manufacturer’s instructions (Guo et al., 2022). 1 μg of total RNA was reverse-transcribed using a first-strand cDNA synthesis kit (Invitrogene). qRT-PCR assays were performed with SYBR Green Mastermix (Bio-Rad) and specific primers for PCR amplification. qRT-PCR data were recorded and analyzed using iQ-PCR (Bio-Rad) equipment and software according to manufacturers’ recommendations (Guo et al., 2022). For each primer pair, the primer efficiency was measured and the melting curve was analyzed. For each experiment, three technical replicates were used. The primers used for the qRT-PCR studies are in Table S1.

## Statistical information

Experiments were usually performed with at least three independent repeats (biological replicates) to ensure the results (Guo et al., 2022; Leng, Yu, et al., 2018). For animal experiments, triplicated breeding was used to obtain statistically significant number of embryos or mice; and statistically significant differences between means of protein levels in the control wildtype and knockout mutants were compared using two-tailed equal-variance independent Student’s t-test (Guo et al., 2022). All other data were determined using a two-tailed equal-variance independent Student’s t-test (Guo et al., 2022). The data in all figures met the assumption of normal distribution for tests. Different data sets were considered to be statistically significant when the P-value was <0.05 (*), 0.01 (**), 0.001 (***), or 0.0001 (****)(Fay & Gerow, 2013).

## Additional files

MDAR checklist

Table-key resources table

## Data Availability

All data generated during this study are included in the manuscript. Uncropped immunoblots, immunostaining, and gel blot images are accessible as source data.

## Acknowledgements

This work was supported by grants from National Institutes of Health (R15CA254827 to HS and R01GM140185 to HZ). The DNA sequencing analysis of animal mutations was supported by the Nevada INBRE Scientific Core Service Award. The flow cytometry analysis was conducted in the Core Facility of Nevada Institute of Personalized Medicine.

## Author Contributions

**Hong Sun** and **Hui Zhang**: Conceptualization, Supervision, Investigation, Writing, Review, Funding acquisition; **Pengfei Guo:** Investigation, writing, Methodology, Animal analysis; **Keshari Rajawasam** and **Tiffany Trinh**: Methodology.

## Appendix

**Appendix Table 1.**
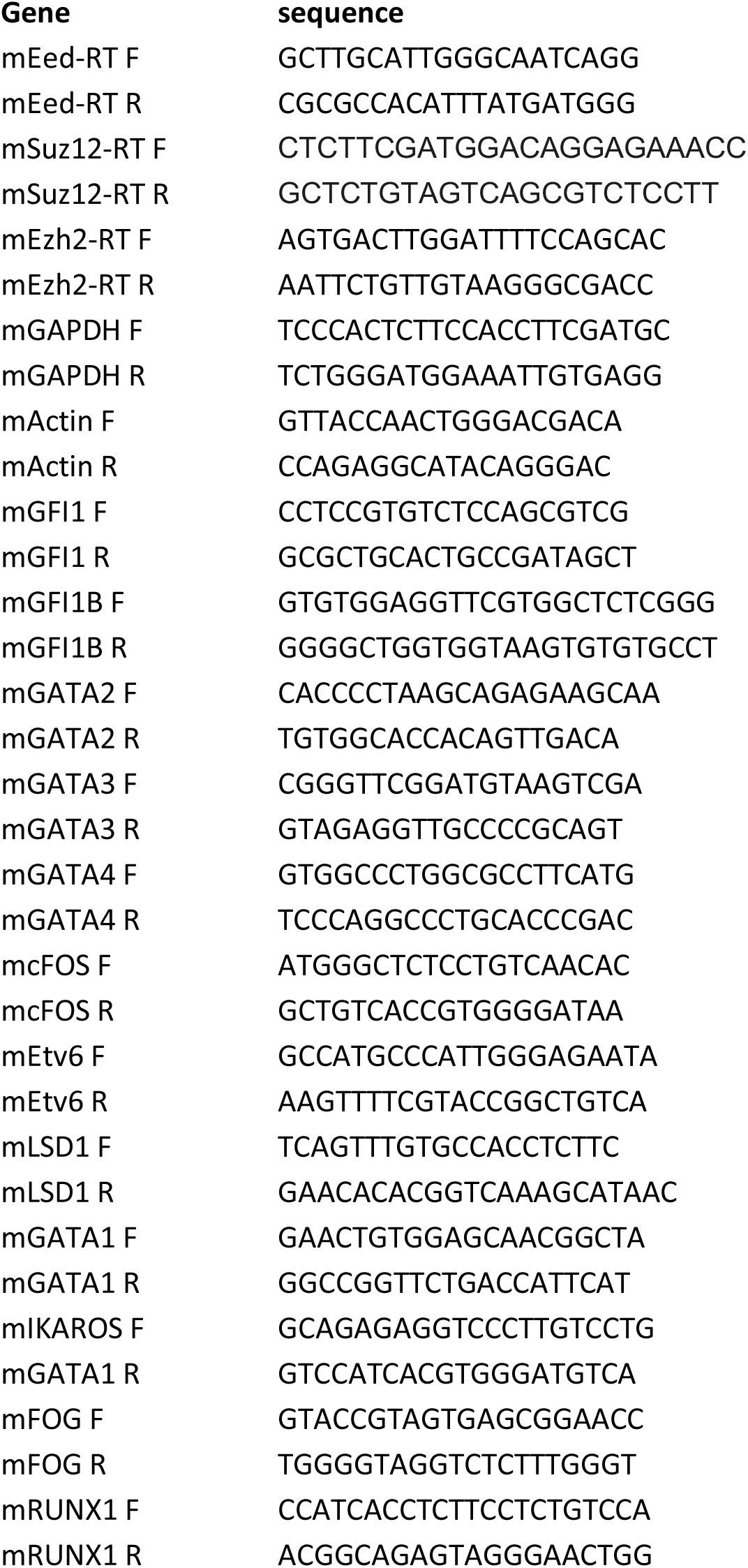
The list of DNA oligonucleotide primers for RT-PCR

---

**Figure 2-figure supplement 1.**
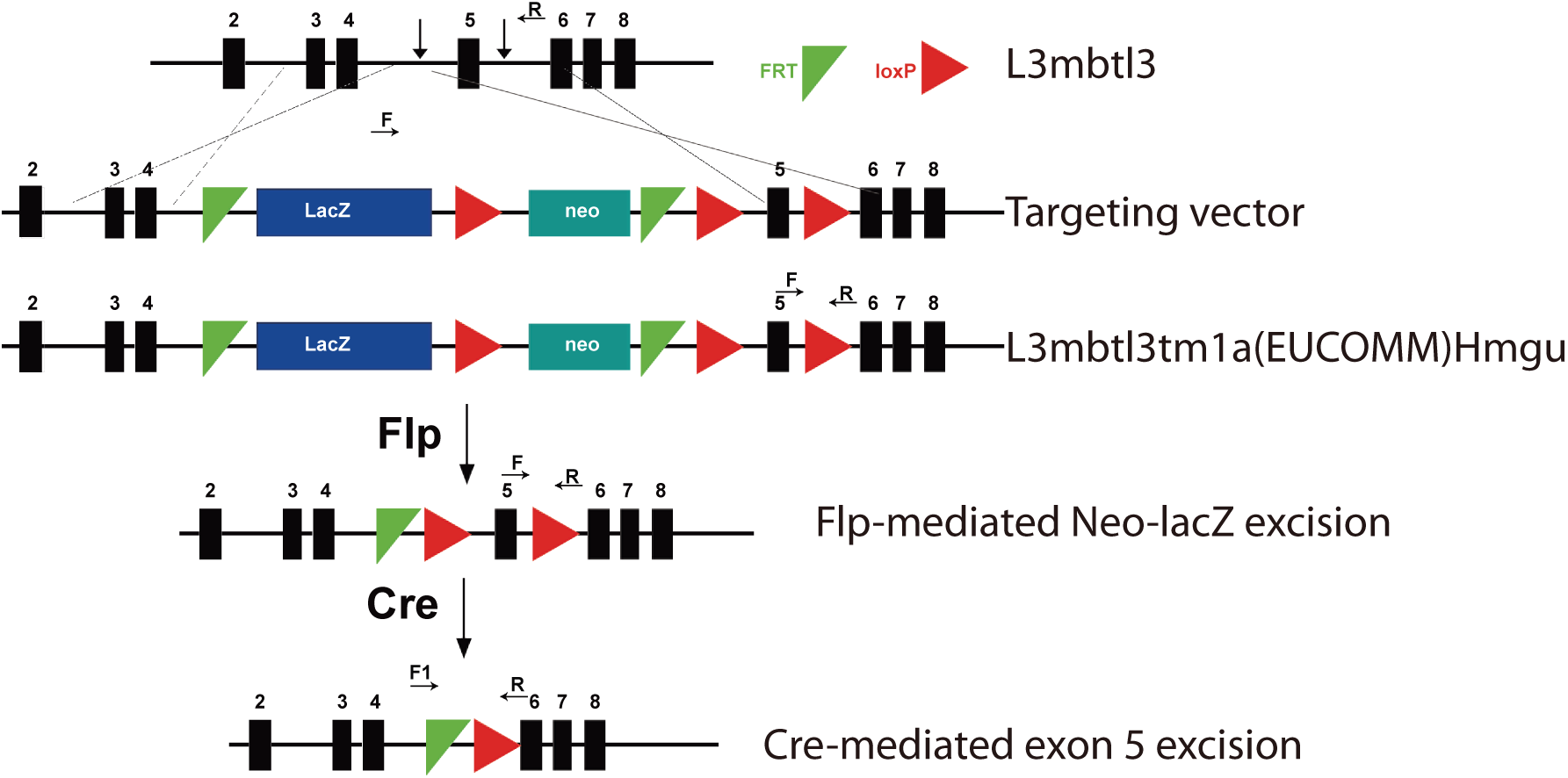
Schematic illustration to generate a conditional flox allele in *L3mbtl3^tm1a(EUCOMM)Hmgu^* with the *FRT-loxP* sites at the exon 5 of the *L3mbtl3* locus. The LacZ-neo fragment was removed by breeding with the *FLPo-10* mice to obtain the conditional *L3mbtl3^flox/+^* mice. Green triangles are *FRT* sites and the red arrows are *loxP* site*s.* F and R are PCR primers for genotyping.

**Figure 4-figure supplement 1.**
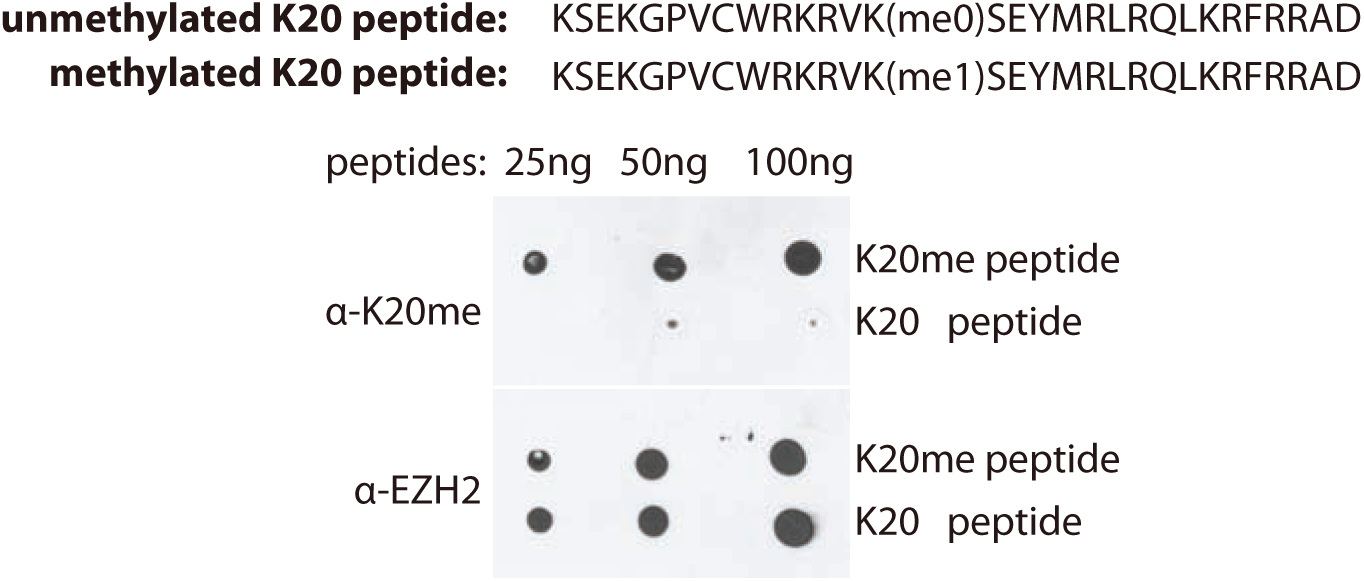
Specificity of anti-methylated K20 peptide antibodies. The unmethylated and monomethylated K20 cognate peptides were spotted onto nitrocellulose membrane. The methylated peptides were detected by the affinity purified anti-mono-methylated K20 peptide and EZH2 antibodies.

**Figure 6-figure supplement 1.**
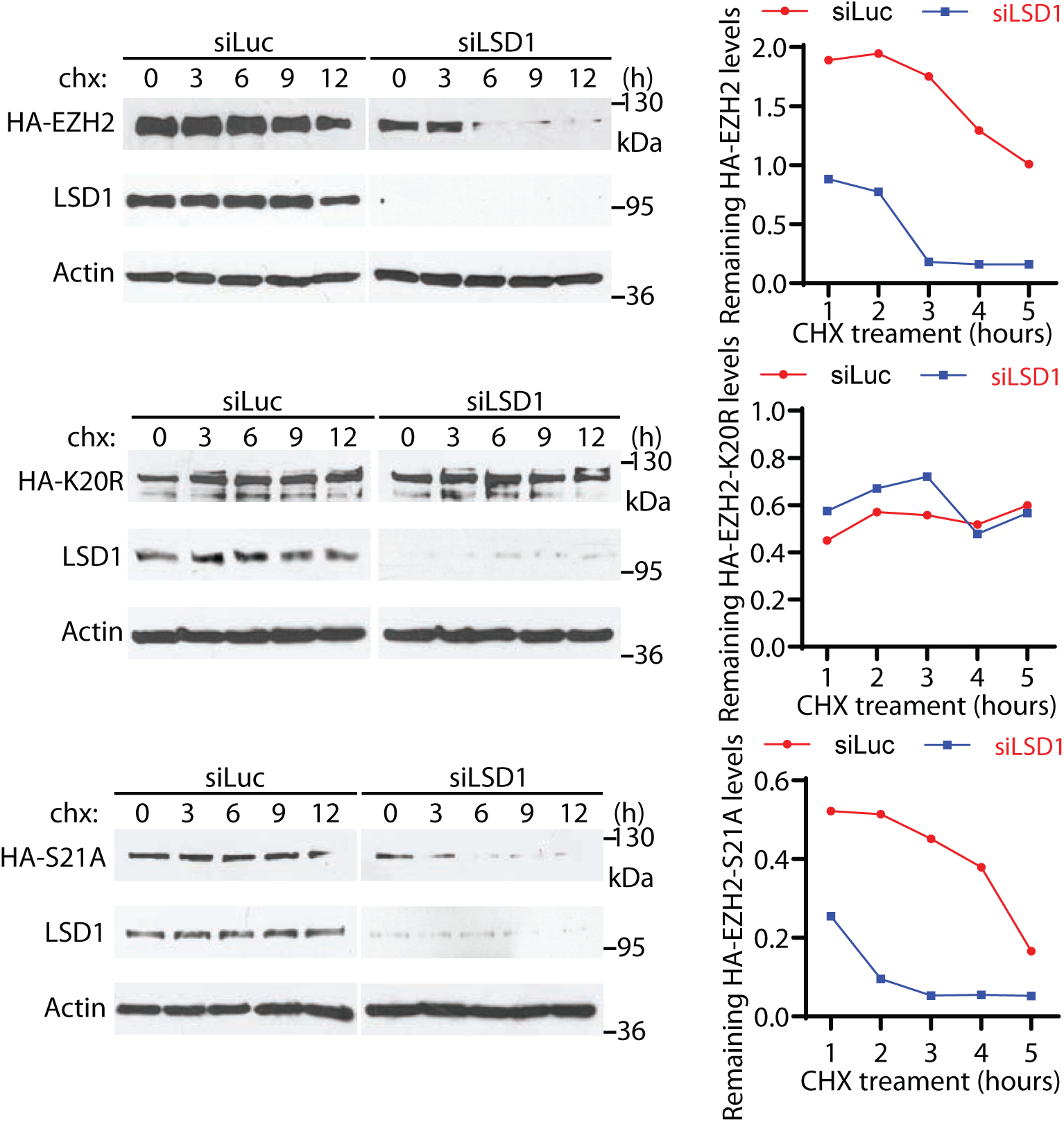
Regulation of EZH2 protein stability by K20 methylation with protein decay assays. The HA-tagged EZH2 WT, K20R, and S21A mutant expressing G401 cells were transfected with 50 nM siRNAs of luciferase or LSD1 for 48 hours, treated with 100 μM cycloheximide (CHX), and collected at the indicated times to measure the half-lives of HA-EZH2 proteins by Western blotting. The protein intensities were quantified by ImageJ software and normalized to the intensity of HA-EZH2 proteins at the zero time when cycloheximide was added.

**Figure 6-figure supplement 2.**
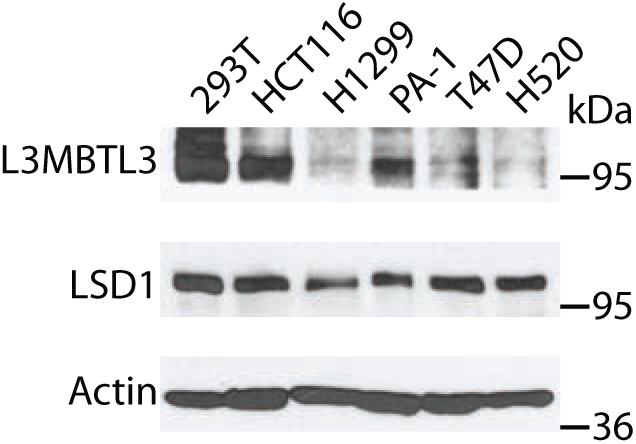
Examination of L3MBTL3 protein levels by West blotting in cancer cell lines.

**Figure 6-figure supplement 3.**
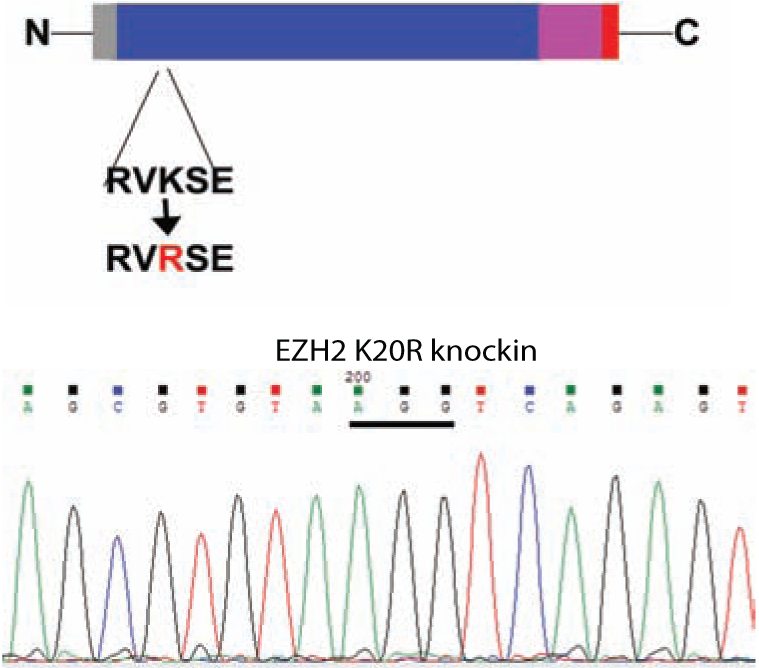
Schematic demonstration of the mouse EZH2 K20R knock-in mutagenesis and a representative DNA sequencing profile performed on the mouse K20R mutant of EZH2.

**Figure 6-figure supplement 4.**
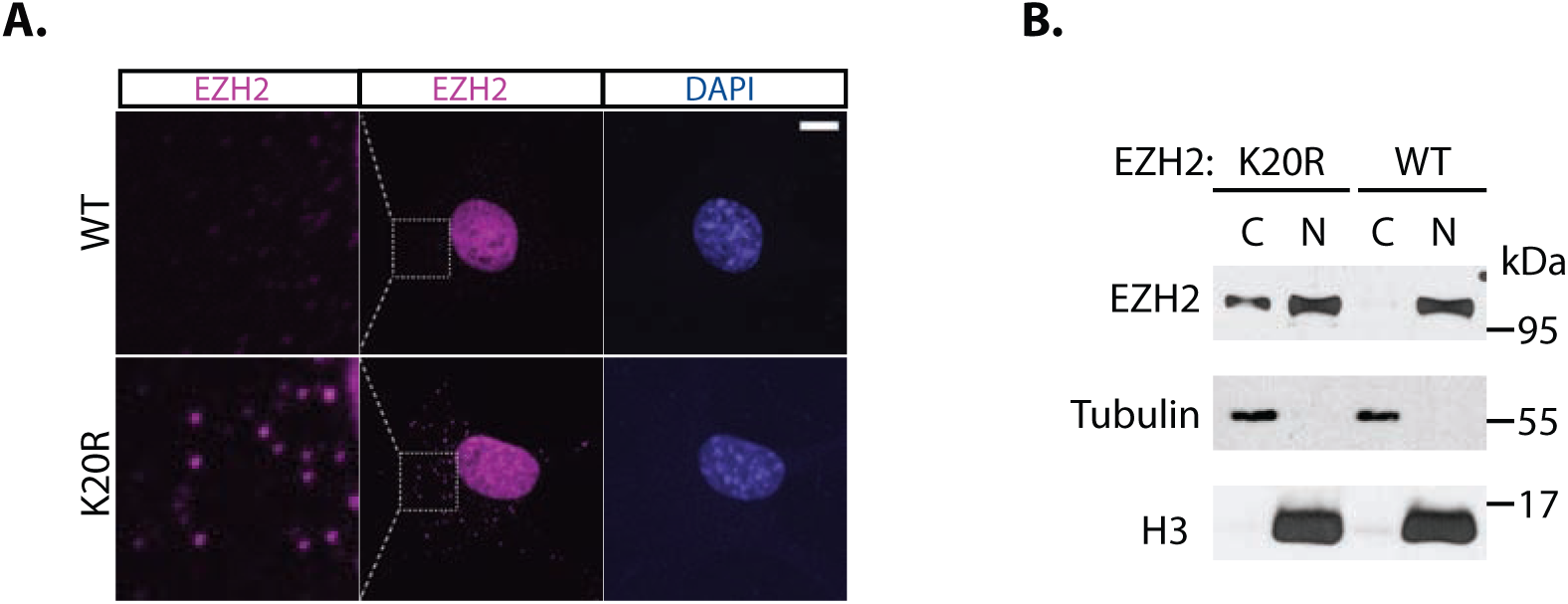
A fraction of K20R mutant protein is in cytoplasm. (**A**) The wildtype and EZH2 K20R MEFs were fixed for anti-EZH2 immunofluorescence staining for EZH2, counter stained with DNA dye, 4′,6-diamidino-2-phenylindole (DAPI). (**B**) Cellular fractionation of the wildtype and EZH2 K20R MEFs into the cytoplasm (C) and nuclear (N) fractions. Proteins were blotted with anti-EZH2, histone H3, and tubulin.

**Figure 7-figure supplement 1.**
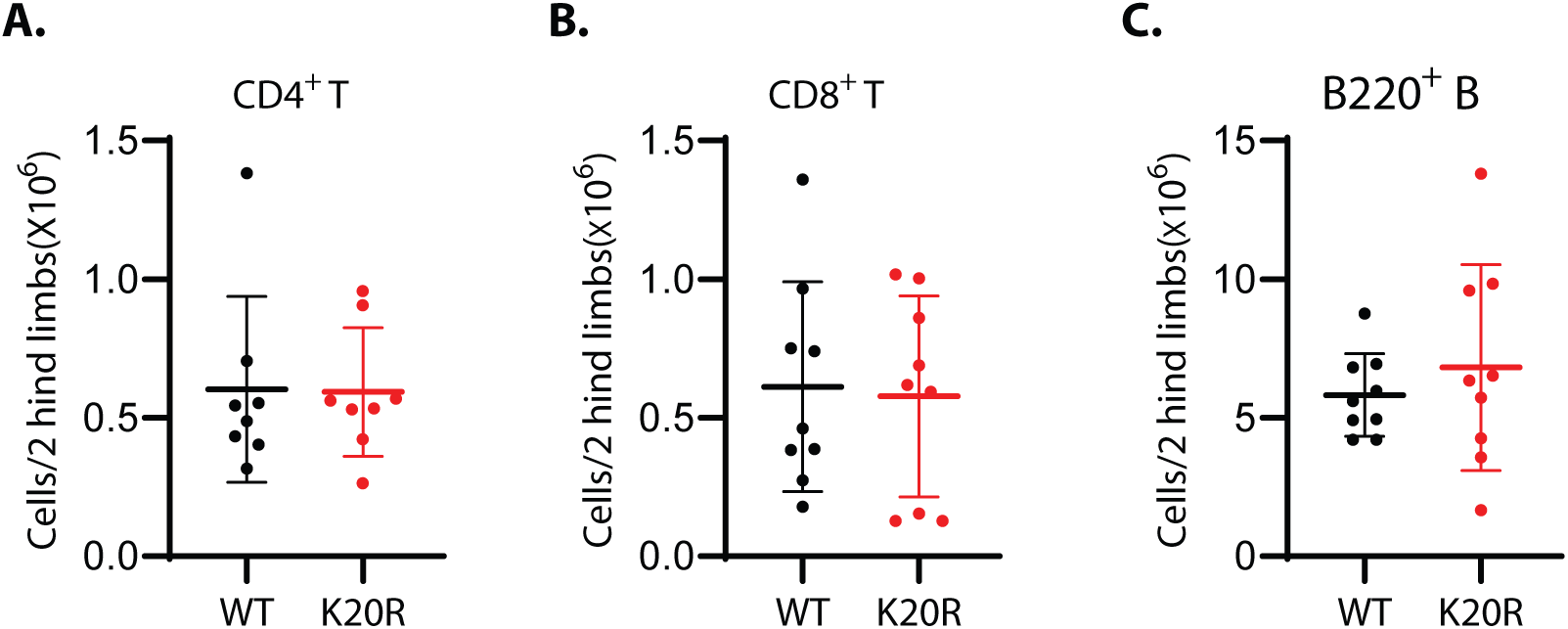
Effects of K20R mutation in T and B cells. (**A-C**) Representative flow cytometric profiles of bone marrow cells harvested from the wildtype control and K20R mutant mice. Flow cytometric plots were gated on the CD4^+^ and CD8^+^ T cells (**A** and **B**) and B cells (**C**, B220+). Values are means ± SEM (n=6∼9). Significance levels were indicated as a two-tailed, unpaired, *t-test*.

